# Sleep and wakefulness in rats: probabilities of state transitions and state-to-state serial correlations in relation to circadian and ultradian phases

**DOI:** 10.1101/2025.03.13.643124

**Authors:** Richard Stephenson

## Abstract

Sleep and wakefulness were recorded from undisturbed male rats (n = 10) for 7 days in a 12:12 h light-dark cycle. Data were partitioned by circadian and ultradian phase before quantification of within-state transition probabilities and between-state serial correlations. Within each joint circadian-ultradian phase, short-term statistical patterns were identified, with quantitative variation between phases. Statistical “signatures” of sleep-wake pattern include positive serial correlations between durations of contiguous states, and between state bouts and inter-bout intervals. There was no correlation between wake bout duration and subsequent sleep episode duration, contradicting an oft-cited assumption of the sleep homeostasis hypothesis. The data confirm a positive correlation between the duration of rapid eye movement (REM) sleep and duration of the subsequent inter-REM interval, which is traditionally interpreted as evidence for a short-term REM homeostat. The present study concludes that the REM homeostat hypothesis is underdetermined. It is suggested that this serial correlation and other sleep-wake patterns can be explained by an alternative non-homeostatic mechanism; transition probability prevailing at the end of a state episode is carried over to the beginning of the next state (“inter-state transition momentum”). The plausibility of this hypothesis was supported by preliminary simulations using a 2nd-order 3-state semi-Markov model.

**Highlights:**

- Wake-related transition probabilities underly circadian and ultradian rhythms.
- Statistical signatures of short-term patterns of sleep-wake state were identified in all circadian and ultradian phases.
- Transition probabilities varied with time in state.
- State transition probabilities were influenced by duration and identity of prior state.
- Sleep homeostasis hypotheses were not supported.
- A hypothesis of state-spanning transition momentum is proposed.

## 1. Introduction

Sleep-wake architecture of polyphasic mammals such as rodents is characterized by frequent transitions between states of wakefulness (WAKE), rapid eye movement sleep (REM) and non-REM sleep (NREM) ^1-8^. State-to-state transitions appear to be probabilistic events and have been described using various quantitative models, many of which assume the Markov property of 1^st^-order dependence wherein probability of the next state transition is dependent only on the current state and is independent of the type and duration of prior states. However, such models are incomplete because it is well known that state transition probabilities can vary with time in state in humans and animals^9-11^, and higher order statistical dependence clearly exists over timescales spanning multiple bouts. For example, serial correlations have been observed in the durations of contiguous sleep-wake bouts ^12, 13^, and a positive serial correlation between REM bout duration and the duration of subsequent inter-REM intervals has also been well documented ^8, 10, 13-15^. It remains unknown how these short-term serial correlations relate to ultradian phase within the light and dark phases of the diurnal cycle.

It was shown recently, in a partial analysis of the dataset used in the present study^16^, that approximately 28% of the amplitude of the diurnal rhythm of wakefulness and sleep is attributable to variation in the durations of the WAKE-dominant (ν_w_) and sleep-dominant (ν_s_) phases of the ultradian cycle. Specifically, the mean ultradian cycle time was significantly shorter in the D phase than in the L phase of the diurnal cycle, owing to a shortened ν_s_ phase that was incompletely offset by a lengthened ν_w_ phase in D (see Table 1 in ^16^). The net result of these diurnal changes in ultradian phase durations was that the fraction of total ultradian cycle time (and total circadian phase time) devoted to the ν_w_ phase (the ultradian duty cycle, D_c_ = ν_w_ / [ν_w_ + ν_s_]) was substantially higher in D than in L. The remaining ∼72% of diurnal rhythm amplitude was found to be attributable to variation in fractional contents of WAKE, NREM and REM within the two phases of the ultradian cycle. The fractional state contents vary as a result of changes in bout duration and bout frequency, which in turn are dependent on the probabilities of state transition. It is presently unknown how these quantitative measures of state dynamics relate to joint circadian - ultradian phase.

The main objective of the present study was to identify quantitative characteristics of sleep-wake architecture in the laboratory rat, and to gain insight into the relevance of short-term statistical patterns of state transition and state bout duration in expression of ultradian and circadian rhythms.

## 2. Methods

### 2.1. Animal maintenance and experimental protocols

The present study is a re-analysis of previously published data and detailed experimental protocols are described elsewhere^6, 16-20^. Ten adult male Sprague-Dawley rats were maintained individually in standard laboratory cages housed in a sound attenuated room at 23 ±1 **°**C, and provided with chow and water *ad libitum* throughout the study. Ten rats (mean ± SEM; body mass 431 ± 15 g at start of recordings) were maintained under a stable 12-h:12-h light-dark cycle for at least 5 weeks before and during the study. They were recorded for 7 full consecutive days, during which they were undisturbed except for a brief interval (usually <5 min) each day immediately after lights on (zeitgeber time, ZT 0 h), at which time the rats and cages were inspected, food and water were replaced, and equipment was checked.

### 2.2. Instrumentation

All animals were instrumented, under isoflurane anaesthesia, with fully implanted biotelemetry devices (model TL10M3-F50-EET, Data Science International, St. Paul, MN) for acquisition of electroencephalogram and electromyogram^21^. Data processing and sleep scoring were performed as described previously^17, 18^. Offline automated sleep scoring was performed using a custom rat sleep autoscoring algorithm written in spreadsheet format (Microsoft Excel, v. 2011, Microsoft Corp., Redmond, WA, USA). This algorithm has been described in detail and validated^17^. Visual analysis of raw data (using sub-samples of EEG and EMG traces) was performed to check the accuracy of the automated scoring system in each rat and all animals used in the study had autoscore versus visual rater concordance >90%. In both automated and visual scoring each 5 s epoch was assigned to one of four categories: wakefulness (WAKE), rapid eye movement sleep (REM) and non-REM sleep (NREM), and artifact (ART). Artifacts were identified algorithmically^18, 22^ and then reclassified by direct visual inspection based on prior and subsequent EEG and EMG patterns.

### 2.3. Data analysis

All analyses were performed using either Microsoft Excel spreadsheets or GraphPad Prism statistical software (version 6.0f for mac OS X, GraphPad Software, San Diego, CA, USA).

Signal to noise enhancement and identification of circadian and ultradian periodicities in the hypnogram is described in detail in a recent paper^16^. Light (L) and dark (D) phases of the circadian cycle were scored directly from the known times of lights on/off. To score ultradian phase, the fractional content of WAKE (F_WAKE_) was recorded after the time series was collapsed into non-overlapping 10 min time bins. This F_WAKE_ time series was smoothed using a 4 h centered moving average, and the original series was zero-reset by subtraction. Ultradian phase transitions were next defined by zero-crossing of a 90 min centered moving average through the zero-reset F_WAKE_ time series. Supra-zero intervals were scored as ν_w_ (WAKE-dominant ultradian phase) and sub-zero intervals were scored as ν_s_ (sleep-dominant ultradian phase).

In each animal, the time series of alternating states was expressed as a list of state bouts and corresponding durations (to the nearest 5 s epoch). The list was partitioned into circadian phase (L and D) and ultradian phase (ν_w_ and ν_s_). Hence, each bout was assigned to one of four joint circadian-ultradian phase categories (Lν_w_, Lν_s_, Dν_w_ and Dν_s_). Analyses were performed separately on the phase-segregated data in each animal.

In this study, a “bout” of each state is defined as any uninterrupted interval (one or more epochs) in a given state of WAKE, NREM or REM. Uninterrupted intervals of sleep consisting of more than one bout (i.e. sequences of NREM and REM) are referred to as “sleep episodes”.

#### 2.3.1. State transition probability versus state bout duration

State transition probabilities were estimated as the fraction of bouts terminating at any given bout duration, t, conditional on the bouts remaining uninterrupted at least until time t. The number of bouts remaining in the sample decreases progressively with increasing bout duration, owing to some of the bouts terminating at each consecutive epoch. Hence, the estimation of true probability by this method becomes correspondingly less precise with increasing bout duration. For this reason, calculations of intra-bout transition probability were performed only for the range of bout durations in which sample sizes were at least 10 bouts.

Conditional probability (per 5 s epoch) of transition of state i to another state j or k (P_im_, where m = j+k) was estimated at a given bout duration, t, as: P_im[t]_ = *n*_[t]_ / *n*_[≥t]_, where, *n*_[t]_ is the number of bouts of duration t, and *n*_[≥t]_ is the number of bouts of duration greater than or equal to t. Conditional probability of state maintenance is the additive inverse: P_ii[t]_ = 1 - P_im[t]_. The probabilities P_im_ and P_ii_ are considered to quantify state instability and state stability, respectively.

As stated, the probability of termination of a bout at time t (i.e. P_im[t]_) is the sum of two transition probabilities: P_im[t]_ = P_ij[t]_ + P_ik[t]_. To calculate these separate post-bout transition probabilities, the relative transition probability was first obtained as follows. For WAKE, P^*^_wr[t]_ = *n*_wr[t]_ / *n*_w[t]_, P^*^_wn[t]_ = *n*_wn[t]_ / *n*_w[t]_, and P^*^_wr[t]_ + P^*^_wn[t]_ = 1, where *n* is number of bouts, first subscript is the initial state (w = WAKE) and second subscript is the subsequent (post-bout) state (n = NREM and r = REM). Likewise for NREM and REM, P^*^_nw[t]_ = *n*_nw[t]_ / *n*_n[t]_, P^*^_nr[t]_ = *n*_nr[t]_ / *n*_n[t]_, P*nw[t] + P*nr[t] = 1, and P*rw[t] = *n*rw[t] / *n*r[t], P*rn[t] = *n*rn[t] / *n*r[t], P*rw[t] +P*rn[t] = 1. Absolute transition probabilities were then estimated for each state by multiplying the total conditional probability of transition, P_im_, by the appropriate relative transition probability, P^*^_ij_ or P^*^_ik_ (P_ij[t]_ = P_im[t]_ x P^*^_ij[t]_).

Thus, in this paper, P_wn_ is the probability of sleep onset (WAKE to NREM transition). P_wr_ is the probability of sleep-onset REM (SOREM; WAKE to REM transition), P_nw_ is the probability of arousal from NREM (NREM to WAKE transition), P_nr_ is the probability of within-sleep REM onset (NREM to REM transition), P_rw_ is the probability of arousal from REM (REM to WAKE transition), and P_rn_ is the probability of within-sleep NREM onset (REM to NREM transition).

Bout-mean probability for any given transition pathway was calculated as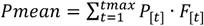 where P_[t]_ is the transition probability at bout duration t epochs, F_[t]_ is the fraction of total bouts with duration t epochs and t_max_ is the maximum bout duration (epochs) in the sample. F_[t]_ sums to 1 across the entire sample.

To evaluate whether the pair of absolute probabilities for any given state varied in parallel over time in state, bouts were segregated into two groups based on bout duration; sub-median and supra-median, and the mean *relative* transition probability was calculated for each group. Paired t-test was used to evaluate significance of differences in relative transition probability between sub-median and supra-median groups in each state, where a significant difference was taken to infer that absolute P_ij_ and P_ik_ varied asynchronously over time in state.

Another way to address this question is to measure state bout durations after each state is segregated according to the identity of subsequent state. If P_ij_ and P_ik_ varied asynchronously over time in state, then shorter and longer bouts will be differentially followed by different states. Thus, states were segregated as follows: WAKE followed by REM (Wr), WAKE followed by NREM (Wn), NREM followed by WAKE (Nw), NREM followed by REM (Nr), REM followed by WAKE (Rw) and REM followed by NREM (Rn). Median bout durations of each segregated state sub-group (i.e. the states denoted by capitalised letters) were recorded in each animal. Significant difference (paired t-test) in median durations of paired groups (i.e. Wr vs Wn, Nw vs Nr and Rw vs Rn) indicates a differential “choice” of post-bout state associated with shorter versus longer bouts. Thus, this categorical analysis was intended to assess whether time in state had an influence on post-bout transition trajectory and served as a confirmatory check on the results of the post-bout relative transition probability analysis described above.

#### 2.3.2. Association between time in state versus identity and duration of prior state

Bouts of each state were segregated according to the identity of preceeding state: WAKE after REM (rW), WAKE after NREM (nW), NREM after WAKE (wN), NREM after REM (rN), REM after WAKE (wR) and REM after NREM (nR). Median bout durations of each segregated state sub-group (i.e. the states denoted by capitalised letters) were recorded in each animal. This categorical analysis was intended to assess whether the identity of prior state had an influence on bout duration of any of the three states. Paired t-test was used to evaluate significance of differences between paired segregated groups in each state (i.e rW vs nW, wN vs rN and wR vs nR).

Spearman rank correlation coefficient (ρ) was used to measure quantitative correlations between the duration of state bouts versus the duration of preceding state bouts for each of the segregated state pairs described above. The threshold for statistical significance (α) was set at 0.025 (Bonferroni correction).

#### 2.3.3. Serial correlations between durations of consecutive bouts within each state

Spearman rank correlation coefficient (ρ) was used to quantify serial correlation between durations of consecutive bouts within each state (i.e. consecutive WAKE bouts, consecutive NREM bouts and consecutive REM bouts) and also between consecutive episodes of sleep. This constitutes a lag 1 autocorrelation of state bout duration, ignoring the intervening inter-bout intervals, and seeks to determine whether there is any evidence of bout-to-bout continuity of state maintenance for a given state.

#### 2.3.4. Serial correlations between bout durations and inter-bout intervals (IBI)

Each bout of a given state is separated by one or more bouts of another state. Thus, each WAKE bout is flanked by preceeding and succeeding inter-WAKE intervals (preIWI and postIWI, respectively). The IWI is, of course, a sleep episode and consists of one or more bouts of NREM and REM. Likewise, each NREM bout is flanked by preceeding and succeeding inter-NREM intervals (preINI and postINI) consisting of one or more bouts of WAKE and REM, and each REM bout is flanked by preceeding and succeeding inter-REM intervals (preIRI and postIRI) consisting of one or more bouts of WAKE and NREM. Thus, the IBI’s are not independent because each bout of each state contributes simultaneously to two IBI’s.

Spearman rank correlation coefficient (ρ) was used to quantify associations between state bout duration versus the durations, cumulative state contents, number of bouts and mean bout durations of the prior (preIBI) and subsequent (postIBI) inter-bout intervals. P values were adjusted by Bonferroni correction.

#### 2.3.5. Sequential association of WAKE sub-types and sleep sub-types

A substantial fraction of all sleep episodes are relatively brief and consist of a single bout of NREM (S_s_). The remainder are longer multi-bout sleep episodes (S_m_) consisting of a variable number of alternating NREM and REM bouts (Fig 1B). It was of interest to determine whether these two types of sleep episode were differentially associated with WAKE duration. Previous reports^2, 4-6, 12, 23, 24^ have suggested, on the basis of various statistical analyses, that WAKE bouts can be sub-divided into two sub-types; a population of “brief” WAKE bouts (W_b_) and a population of “long” WAKE bouts (W_l_). In this study, a threshold duration (T_bl_) was used to segregate WAKE bouts into the two sub-types in each circadian-ultradian phase as follows.

**Fig. 1.**
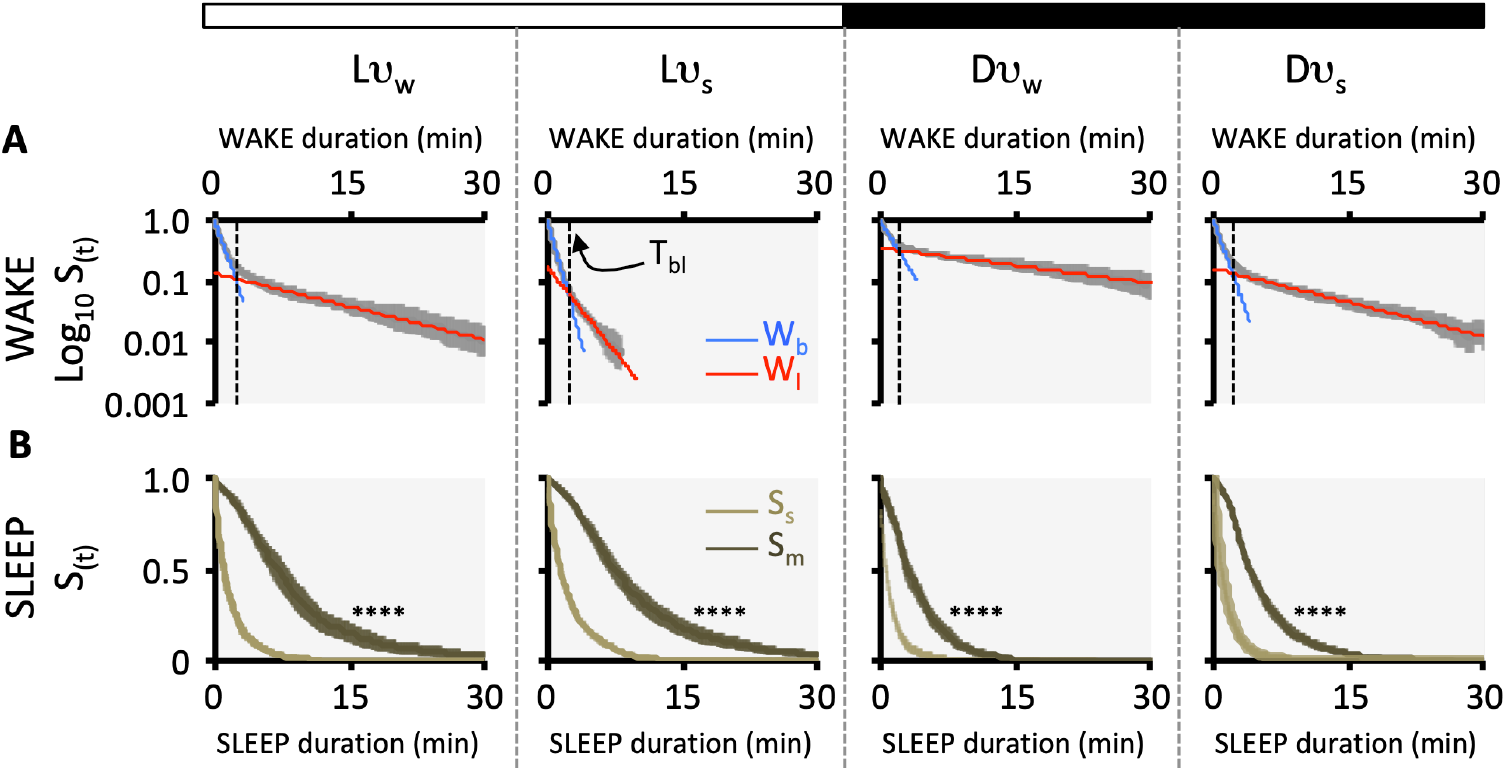
Mean (± 95% CI, n = 10) survival functions (S_(t)_) versus state duration (min) for WAKE bouts (**A**) and SLEEP episodes (**B**). Light and dark phases of the 12:12 h photocycle are indicated schematically as white (L) and black (D) bars at top. Ultradian rhythms (WAKE-dominant ν_w_, and sleep-dominant ν_s_ phases) were present in both L and D and data were subdivided into each of four joint circadian-ultradian phases (Lν_w_, Lν_s_, Dν_w_ and Dν_s_). **A**. WAKE bout survival curves are presented on a semi-logarithmic scale to illustrate the method by which bouts were segregated into “brief” WAKE (W_b_) and “long” WAKE (W_l_) bout sub-types. A threshold duration (T_bl_, dotted vertical line) was defined as the duration at which linear regression lines for W_b_ (blue) and W_l_ (red) intersect. **B**. Sleep episodes were segregated on the basis of the number of bouts per episode. Sleep episodes containing a single bout of NREM or (rarely) REM (S_s_, light olive), were contrasted with multi-bout sleep episodes (S_m_, dark olive). Sleep episode survival curves are presented on linear coordinates to emphasize the difference in durations of the two sleep sub-types (log-rank test; ****, P < 0.0001).

Kaplan-Meier curves were computed in the standard way in each animal:

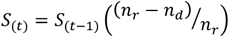

where S_(t)_ is surviving fraction in the epoch ending at time t, S_(t-1)_ is surviving fraction at the end of the immediately preceding epoch, *n*_r_ is the number of bouts remaining at the start of epoch (t), and *n*_d_ is the number of bouts that terminate during epoch (t). A bi-exponential function was fit to the WAKE bout survival curve:

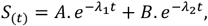

where, A is the proportion of the “fast” decaying component (subscript 1), B is the proportion of the “slow” decaying component (subscript 2) in the curve, and λ is a scale parameter. Specifically, the parameter λ is the exponential rate constant and the inverse of λ is the exponential time constant (mean bout lifetime, τ; 1/e = 0.368 bouts survive), 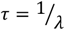.

The threshold bout duration (T_bl_) separating the two WAKE sub-types was obtained using an “exponential peeling” approach^25^. Log S_(t)_ was plotted against t and the linear right tail was fitted by least-squares linear regression. The fitted line was then extrapolated back to the ordinate (i.e. at t = 0). This regression yielded λ_2_ (slope), B (Y intercept) and A (1 - B). The antilogs of the fitted values of log S_(t)_ versus t were then subtracted from the original S_(t)_ values and the resulting differences were again log-transformed before fitting a new linear regression. This regression yielded λ_1_. Finally, the time of demarcation between “brief” and “long” sub-types of WAKE bout (T_bl_), was defined as the time of intersection of the two regression lines (see Fig 1A). Comparing the proportions of WAKE bouts assigned to “brief” and “long” sub-types using this threshold, versus the proportions determined by nonlinear regression of survival curves, group mean error (± 95% CI, n=10) due to using the threshold was estimated to be Lν_w_, 3.6 ±1.9 %; Lν_s_, 0.4 ± 0.3 %; Dν_w_, 2.0 ± 5.3 %, Dν_s_, 1.7 ± 4.7%.

For each of the circadian-ultradian phases, the numbers of bouts of each WAKE sub-type (W_b_, W_l_) and the numbers of episodes of each sleep sub-type (S_s_, S_m_) were recorded, together with the numbers of occurrences of the 8 possible 2-state sequences (“state pairs”). State pairs were defined for the four WAKE to sleep sequences W_b_-S_s_, W_b_-S_m_, W_l_-S_s_, and W_l_-S_m_, and for the four sleep to WAKE sequences S_s_-W_b_, S_s_-W_l_, S_m_-W_b_, and S_m_-W_l_. The objective was to determine whether the sleep sub-types were paired randomly with the WAKE sub-types (the null hypothesis of statistical independence) or whether some sequences occurred preferentially in any of the circadian-ultradian phases.

The null hypothesis was tested in each animal individually using χ^2^ tests: χ^2^ = Σ([observed – expected]^2^ / expected). For each of the sets of state pairs (WAKE - sleep and sleep - WAKE), data were entered into a 2 x 2 contingency table and expected values were calculated from marginal values as (*n*_p_ x *n*_.q_)/*n*, where *n*_p_ is the row sum (i.e. the total number of bouts of the first state of each pair), *n*_.q_ is the column sum (i.e. the total number of bouts of the second state of each pair) and *n* is the total matrix sum.

Percent deviation from expected values was obtained for each pairing sequence (using observed and expected values in each cell of the matrix) and grouped across animals before applying a one-sample t-test to determine the probability that the deviation of group data differs from the null hypothesis of zero deviation.

## 3. Results

### 3.1. Transition probabilities versus time in state

Estimates of transition probabilites (absolute, per 5 s epoch) over time in state are shown in Fig 2. The group mean curves are illustrated in bold colour to highlight general trends (WAKE, blue; NREM, green; REM, red) and individual animal data are shown in fine line plots to highlight variability. All probabilities varied with time in state in all of the circadian-ultradian phases. Qualitative features common to all phases and states were an initial interval of steady or decreasing probability, followed by a linear or monotonic rise. Despite this similarity in overall pattern, the rates of change, as well as the bout-mean probability can be seen to differ both between states and across the four circadian-ultradian phases.

**Fig. 2.**
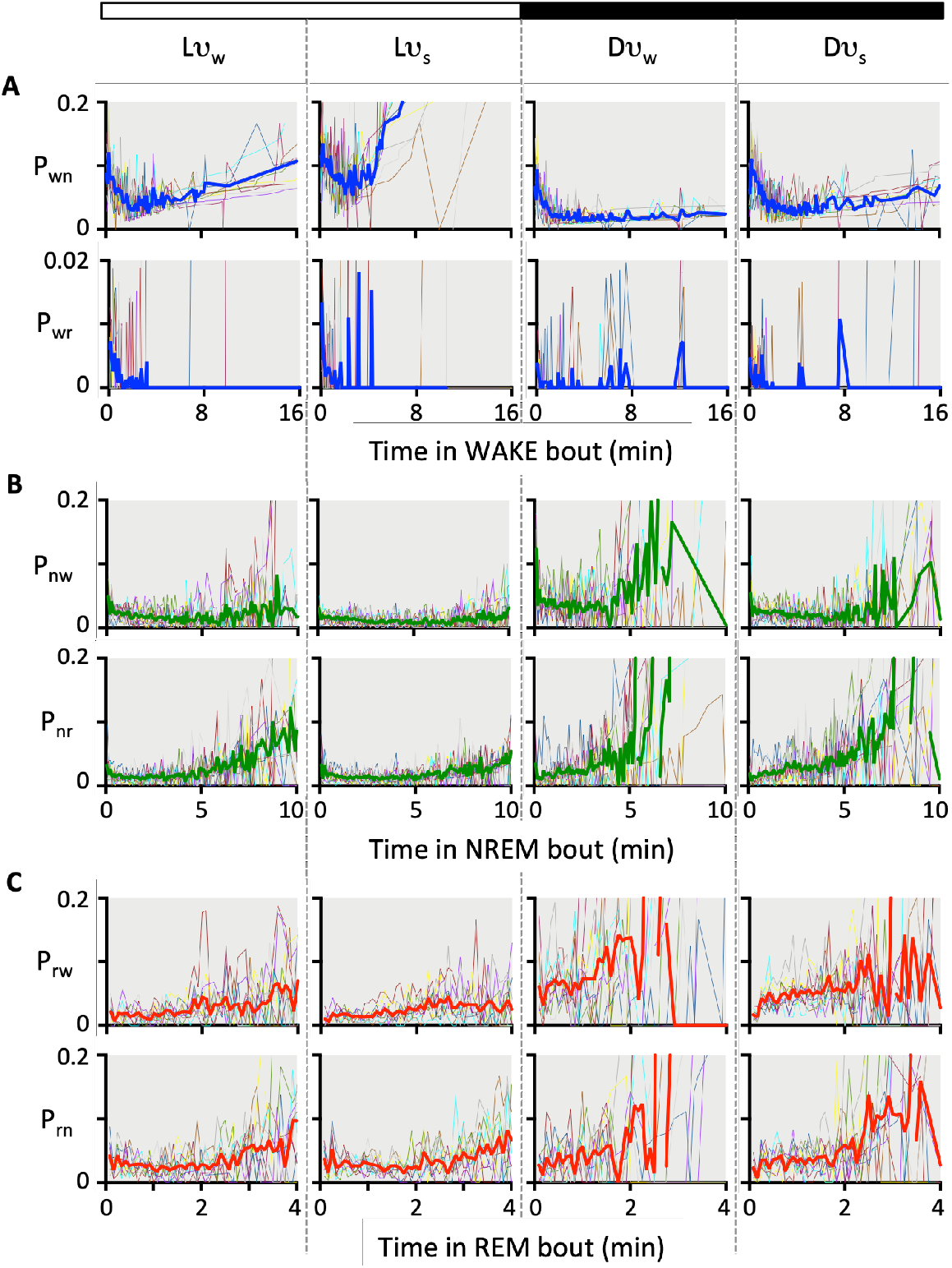
State transition probabilities (per 5 s epoch) versus time in state (min) for WAKE (**A**, blue), NREM (**B**, green) and REM (**C**, red). Each row of panels depict individual animal data (fine lines) and group mean data (thick coloured lines) for a single transition pathway in each of four joint circadian-ultradian phases (Lν_w_, Lν_s_, Dν_w_ and Dν_s_). Each state can transition to one of two other states, and the sum of the two transition probabilities is a measure total state instability. The converse - state stability - is quantified as the state maintenance probability, which is the additive inverse of total state transition probability (i.e. P_ii_ = 1 - (P_ij_ + P_ik_)). Estimates of transition probability become progressively less precise with increasing time in state owing to progressively smaller sample sizes, and this is reflected in significant heteroscedasticity in each plot. Each transition is characterised by time-varying probability in all four circadian-ultradian phases. **A**. WAKE bouts transition to either NREM or REM with respective probabilites P_wn_ and P_wr_. Sleep-onset REM (SOREM’s) are very infrequent and the magnitude of P_wr_ is correspondingly lower (as indicated by the 10x smaller scale for P_wr_). Since P_wn_ >> P_wr_, WAKE maintenance P_ww_ ≈ 1 - P_wn_. Note that P_wn_ (and hence P_ww_) are strongly related to circadian-ultradian phase. **B**. NREM bouts transition to either WAKE (arousal) or REM (sleep cycling) with respective probabilites P_nw_ and P_nr_. Note that the timecourses of P_nw_ and P_nr_, although similar in their overall pattern, are not synchronous so that the relative probabilities of arousal and sleep cycling vary as a function of time in NREM (i.e. relative likelihood of REM onset increases progressively during NREM). **C**. REM bouts transition to either WAKE (arousal) or NREM (sleep cycling) with respective probabilites P_rw_ and P_rn_. Note that within each circadian-ultradian phase, the timecourses of P_rw_ and P_rn_, were similar in their overall pattern and varied approximately synchronously across a REM bout so that the relative probabilities of arousal and sleep cycling did not change consistently as a function of time in REM.

The overall contribution of the various state transitions to the expression of circadian and ultradian rhythms of wakefulness and sleep are summarized using the bout-mean probability data and are shown in Fig 3.

**Fig. 3.**
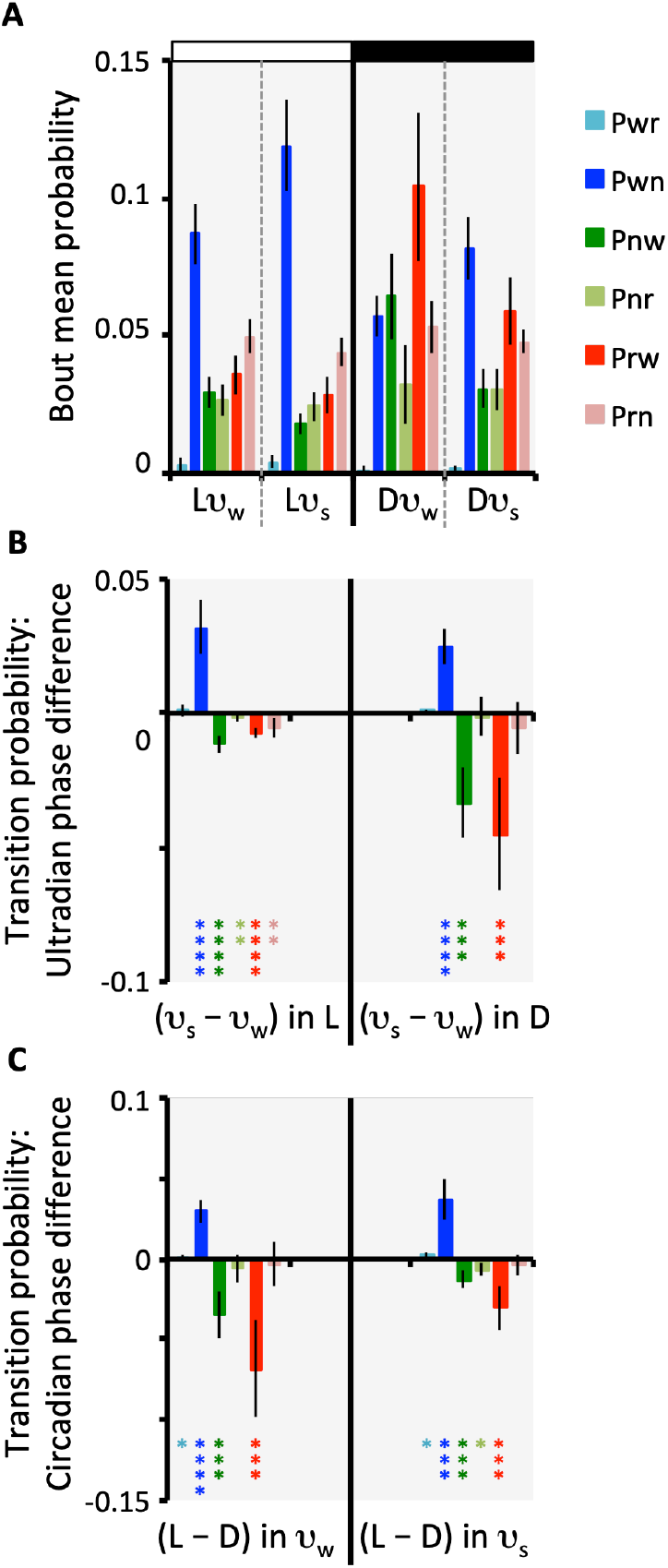
**A**. Bout-mean transition probability and its relation to joint circadian-ultradian phase. These data express the overall average probability values in each of the three states of WAKE, NREM and REM. **B**. Ultradian phase differences (ν_s_ - ν_w_) in each of the L and D circadian phases. Here the data shown in panel **A** are re-expressed as ultradian phase differences to estimate the extent to which each of the transition probabilities vary as a function of ultradian phase. **C**. Circadian phase differences (L - D) in each of the WAKE-dominant (ν_w_) and sleep-dominant (ν_s_) ultradian phases. Here the data shown in panel **A** were re-expressed as circadian phase differences to estimate the extent to which each of the transition probabilities vary as a function of circadian phase. Data in all panels are shown as mean (± 95% CI, n = 10), and statistical significance was determined using paired t-test: *, P < 0.05; **, P < 0.01; ***, P < 0.001; ****, P < 0.0001.

#### 3.1.1. WAKE

P_wr_ was extremely low in all four circadian-ultradian phases (note the expanded scale in Fig 2A), and WAKE to REM transitions constituted only approximately 2.5% of 1360 ± 193 sleep onsets per animal. There were no detectable ultradian differences in bout-mean P_wr_ (Fig 3B), but there was a small statistically significant circadian rhythm - an increase in P_wr_ in the L phase compared with the D phase. This circadian difference was present in both ν_w_ and ν_s_ ultradian phases (Fig 3C).

P_wn_, the probability of WAKE to NREM transition, was relatively high at the start of WAKE bouts (Fig 2A) and decreased monotonically to a minimum after approximately 2-4 min of wakefulness in all four circadian-ultradian phases. P_wn_ subsequently increased at rates that differed between circadian-ultradian phases. To compare across circadian-ultradian phases, bout-mean P_wn_ was used as an estimate of the overall magnitude of the intra-bout probability timecourse (Fig 3A). The ν_s_ - ν_w_ difference in mean P_wn_ was obtained in each of the circadian L and D phases in order to estimate the respective ultradian ranges (Fig 3B). In addition, the L - D difference in mean P_wn_ was obtained for each of the ultradian ν_w_ and ν_s_ phases in order to estimate the respective circadian ranges (Fig 3C). It can be seen in Fig 3 that variation in bout-mean P_wn_ contributed significantly to both ultradian and circadian rhythms. A significant ultradian difference in bout-mean P_wn_ was observed in both the daytime (L) and at night (D) (Fig 3A,B). Furthermore, the circadian rhythm in bout-mean P_wn_ was present in both of the ultradian phases (Fig 3A,C).

In Fig 2A, the change in P_wn_ with time in state appears to differ from that of P_wr_, although the validity of the comparison is open to question owing to small sample sizes for P_wr_. A difference in the timecourses of the two probabilities would be expected to bias the post-bout transition trajectory as a function of WAKE bout duration. Fig 2A suggests that for WAKE bouts longer than approximately 2 min duration, P_wn_ tended to rise gradually (especially during the L phase), whereas P_wr_ remained close to zero. Hence, SOREM’s are expected to mainly follow shorter WAKE bouts. The relative probability of post-WAKE transition trajectory (P^*^_wn_) was not normally distributed across animals and so the statistical significance of differences in P^*^_wn_ between short (sub-median) WAKE bouts and long (supra-median) WAKE bouts were tested using Dunn’s multiple comparisons test. This test identified a difference (P_adj_ = 0.0283) only in the Lν_s_ circadian-ultradian phase.

#### 3.1.2. NREM

P_nw_ (probability of arousal) and P_nr_ (probability of REM onset) both varied over the course of a NREM bout (Fig 2B). The timecourses of the two transition probabilities in NREM followed a gradual decline over the first approximately 5 min of the bout indicating an initial progressive stabilisation of the NREM state. This was more obvious in the L phase than in D. Following this intial stage, both transition probabilities gradually increased, indicating an increasing likelihood of bout termination. The latter was especially marked in the D phase of the circadian cycle and in the ν_w_ phase of the ultradian cycle.

Although the general trends described above were similar in P_nw_ and P_nr_, these two transition probabilities did not vary in exact synchrony, giving rise to progressive variation in post-bout transition trajectory. Specifically, the rise in P_nr_ tended to preceed and exceed, that of P_nw_ (Fig 2B), indicating an increasing relative likelihood of REM onset after longer NREM bouts. This was the case in all circadian-ultradian phases, and was confirmed by two observations: median bout durations of NREM followed by REM (Nr) were significantly longer than median bout durations of NREM followed by arousal (Nw) (Holm-Sidak’s multiple comparisons test: Lν_w_, P_adj_ = 0.0011; Lν_s_, P_adj_ = 0.0042; Dν_w_, P_adj_ = 0.0006; Dν_s_, P_adj_ = 0.0005), and the relative probability of post-NREM arousal (P^*^_nw_) was statistically significantly higher for sub-median NREM bouts than for supra-median NREM bouts (Holm-Sidak’s multiple comparisons test: Lν_w_, P_adj_ = 0.0020; Lν_s_, P_adj_ = 0.0024; Dν_w_, P_adj_ = 0.0003; Dν_s_, P_adj_ = 0.0008).

Bout-mean values were used to assess the dependences of NREM transition probabilities (P_nw_ and P_nr_) on circadian and ultradian phases (Fig 3A). Statistically significant ultradian rhythms were detected in both L and D for bout-mean P_nw_, whereas a very small amplitude ultradian rhythm in P_nr_ was present only in L (Fig 3B). Circadian rhythms were present in P_nw_ during both the ν_w_ and ν_s_ ultradian phases (Fig 3C). However bout-mean P_nr_ exhibited only very small L vs D differences, and this was statistically significant only in the ν_s_ ultradian phase (Fig 3C). Thus, bout-mean P_nw_ varied considerably across both ultradian and circadian rhythms, whereas bout-mean P_nr_ was minimally dependent on ultradian-circadian phase.

#### 3.1.3. REM

SOREM’s were infrequent in all animals (34 ± 21 bouts per animal over the week of recordings). SOREM bouts represented <4% of the 908 ± 153 REM bouts recorded per animal.

Probabilities of REM transition to NREM (P_rn_) or arousal (P_rw_) both varied over time during REM bouts, in all four circadian-ultradian phases (Fig 2C). The timecourse featured an initial interval of stability (steady or declining transition probability) for the first 1-2 min, followed by a trend for increasing transition probability as time in REM progressed further.

The timecourses of P_rw_ and P_rn_ were generally quite similar to each other (Fig 2C); that is, the two probabilities changed essentially in synchrony over the course of a REM bout. This was confirmed by absence of significant differences between median durations of REM bouts that ended in arousal (Rw) compared with those that were followed by NREM (Rn), and by absence of significant differences between the relative probability of post-REM transition trajectory (P^*^_rw_) in sub-median REM bouts versus supra-median REM bouts.

### 3.2. Categorical association between state duration and identity of prior state

The median bout duration of WAKE immediately after REM was marginally longer than that of WAKE after NREM, but this was statistically significant only in the D phase (rW > nW; paired t-test: Lν_w_, P = 0.776; Lν_s_, P = 0.105; Dν_w_, P = 0.034; Dν_s_, P = 0.014).

The association between median NREM bout duration and identity of prior state was of small magnitude and was statistically significant only during the Dν_w_ circadian-ultradian phase (rN > wN; paired t-test: Lν_w_, P = 0.923; Lν_s_, P = 0.051; Dν_w_, P = 0.002; Dν_s_, P = 0.485).

There were statistically significant differences between median bout durations for REM after WAKE versus REM after NREM only during the Lν_w_ and Lν_s_ circadian-ultradian phases (wR > nR; paired t-test: Lν_w_, P = 0.009; Lν_s_, P < 0.0001; Dν_w_, P = 0.057; Dν_s_, P = 0.342), although again the small sample size for wR casts doubt on the reliability of this result.

### 3.3. Quantitative association between state duration and duration of prior state

In NREM-WAKE pairs (nW), the bout duration of WAKE was not correlated with the duration of prior NREM bouts in any circadian-ultradian phase (Fig 4). By contrast, WAKE bout duration was found to be positively correlated with preceeding REM bout duration in REM-WAKE pairs (rW) (Fig 4). This was most apparent in the ν_s_ ultradian phases of both L and D (i.e. Lν_s_ and Dν_s_), and was also present in Dν_w_ (but not in Lν_w_).

**Fig. 4.**
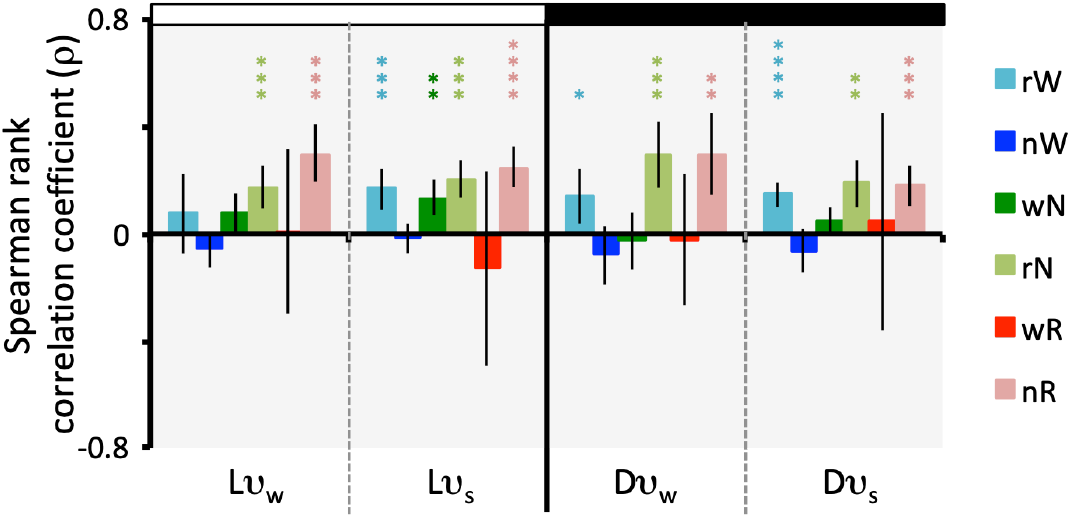
Spearman rank correlation coefficient (ρ; mean ± 95% CI, n = 10) for bout durations of serial pairs of states: REM-WAKE (rW), NREM-WAKE (nW), WAKE-NREM (wN), REM-NREM (rN), WAKE-REM (wR) and NREM-REM (nR). Data were analyzed separately for all four circadian-ultradian phases. One-sample t-test was used to determine statistical significance of group correlation coefficients: *, P < 0.05; **, P < 0.01; ***, P < 0.001; ****, P < 0.0001.

In WAKE-NREM pairs (wN), the duration of NREM bouts was weakly positively correlated with duration of prior WAKE bouts only in the Lν_s_ circadian-ultradian phase (Fig 4). There was no significant correlation in any other phase. By contrast, a significant positive correlation was detected between NREM bout duration and preceeding REM bout duration (rN) in all four circadian-ultradian phases (Fig 4).

There was no detectable correlation between REM bout duration and prior WAKE bout duration in WAKE-REM pairs (wR) in any circadian-ultradian phase (Fig 4). However, ρ coefficients varied greatly between animals owing to very small sample sizes for SOREM bout pairs. By contrast, REM bout durations were significantly positively correlated with durations of preceeding NREM bouts in NREM-REM pairs (nR) in all four circadian-ultradian phases (Fig 4).

### 3.4. Serial correlations between consecutive bouts within each state

There were significant positive correlations between durations of consecutive episodes of sleep, between consecutive bouts of WAKE and between consecutive bouts of NREM in all four circadian-ultradian phases (Fig 5). This suggests a statistical “persistence” of state maintenance from one bout to the next bout of the same state. That is, this serial correlation quantifies a statistical tendency for short bouts and long bouts to each occur in trains of two or more. In contrast, consecutive bouts of REM were not correlated in any circadian-ultradian phase.

**Fig. 5.**
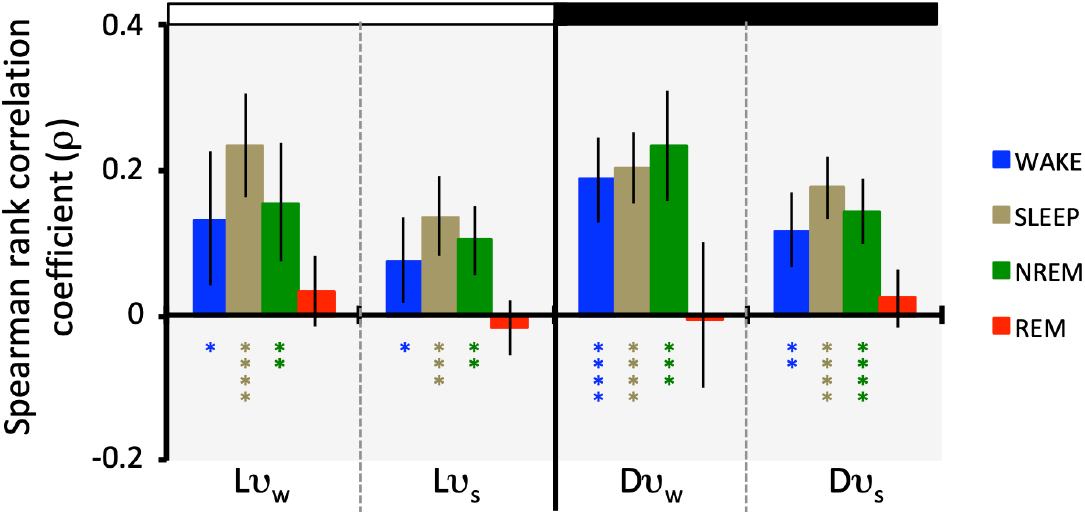
Spearman rank correlation coefficient (ρ; mean ± 95% CI, n = 10) for bout durations of consecutive bouts of a given state. Data were analyzed separately for all four circadian-ultradian phases. One-sample t-test was used to determine statistical significance of group correlation coefficients: *, P < 0.05; **, P < 0.01; ***, P < 0.001; ****, P < 0.0001

### 3.5. Serial correlations between state bout durations and inter-bout intervals

#### 3.5.1. WAKE bout durations versus inter-WAKE intervals (IWI)

WAKE bout duration was weakly, but significantly, positively correlated with the duration of preIWI (i.e. prior sleep episodes) in the Lν_s_ phase only (Fig 6A). In this phase, WAKE bout duration was positively correlated with preIWI cumulative time in REM (Fig 6C), preIWI bout count (Fig 7A) and mean REM bout duration (Fig 7C). In the Dν_w_ and Dν_s_ circadian-ultradian phases, WAKE bout duration was significantly positively correlated with preIWI mean REM bout duration, but in these phases there was no signifcant correlation with preIWI bout count (Fig 7A). Furthermore, in Dν_w_ and Dν_s_ there was a trend for a negative correlation between WAKE bout duration and preIWI cumulative NREM (Fig 6B) and preIWI mean NREM bout duration (Fig 7B) that negated the aforementioned positive correlations with REM, resulting in no net correlation with preIWI duration (Fig 6A).

**Fig. 6.**
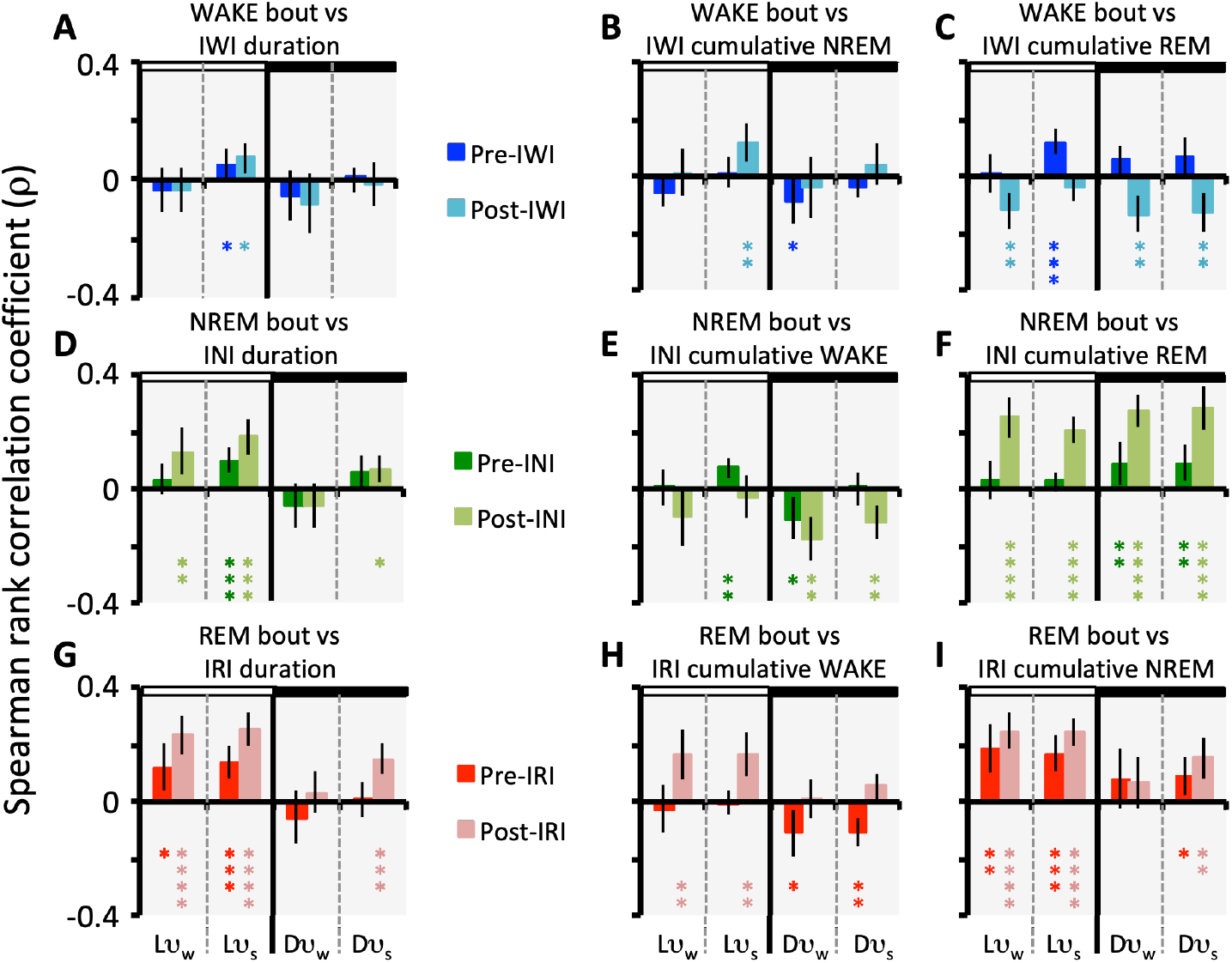
Correlation analysis of bout duration versus durations and state contents of prior (pre) and subsequent (post) inter-bout intervals in all four circadian-ultradian phases. Spearman rank correlation coefficient (ρ) was used in all cases, using average rank for ties. **A**. WAKE bout duration versus durations of pre-(dark blue) and post-(light blue) inter-wake intervals (IWI). **B**. WAKE bout duration versus pre- and post-inter-wake interval cumulative NREM content. **C**. WAKE bout duration versus pre- and post-inter-wake interval cumulative REM content. **D**. NREM bout duration versus durations of pre-(dark green) and post-(light green) inter-NREM intervals (INI). **E**. NREM bout duration versus pre- and post-inter-NREM interval cumulative WAKE content. **F**. NREM bout duration versus pre- and post-inter-NREM interval cumulative REM content. **G**. REM bout duration versus durations of pre-(dark red) and post-(light red) inter-REM intervals (IRI). **H**. REM bout duration versus pre- and post-inter-REM interval cumulative WAKE content. **I**. REM bout duration versus pre- and post-inter-REM interval cumulative NREM content. Data are mean ± 95% CI, n = 10. Statistical significance was determined by one-sample t-test for group (n=10) correlation coefficients: *, P < 0.05; **, P < 0.01; ***, P < 0.001; ****, P < 0.0001.

**Fig. 7.**
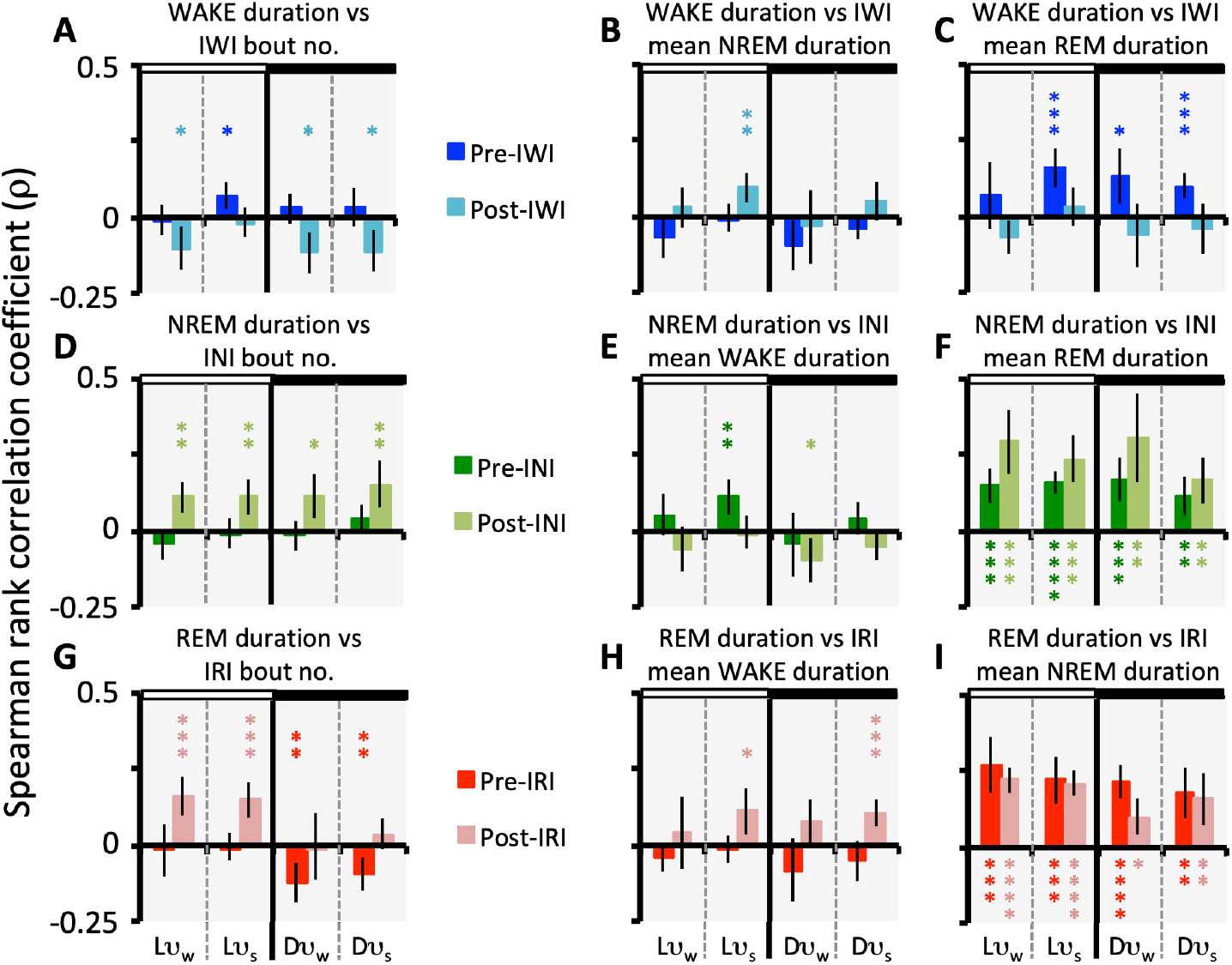
Correlation analysis of bout duration versus numbers of bouts and mean bout durations within prior (pre) and subsequent (post) inter-bout intervals in all four circadian-ultradian phases. Spearman rank correlation coefficient (ρ) was used in all cases, using average rank for ties. **A**. WAKE bout duration versus numbers of bouts in pre-(dark blue) and post-(light blue) inter-wake intervals (IWI). **B**. WAKE bout duration versus mean NREM bout duration in pre- and post-inter-wake intervals. **C**. WAKE bout duration versus mean REM bout duration in pre- and post-inter-wake intervals. **D**. NREM bout duration versus numbers of bouts in pre-(dark green) and post-(light green) inter-NREM intervals (INI). **E**. NREM bout duration versus mean WAKE bout duration in pre- and post-inter-NREM intervals. **F**. NREM bout duration versus mean REM bout duration in pre- and post-inter-NREM intervals. **G**. REM bout duration versus numbers of bouts in pre-(dark red) and post-(light red) inter-REM intervals (IRI). **H**. REM bout duration versus mean WAKE bout duration in pre- and post-inter-REM intervals. **I**. REM bout duration versus mean NREM bout duration in pre- and post-inter-REM intervals. Data are mean ± 95% CI, n = 10. Statistical significance was determined by one-sample t-test for group (n=10) correlation coefficients: *, P < 0.05; **, P < 0.01; ***, P < 0.001; ****, P < 0.0001.

WAKE bout duration was weakly, but significantly, positively correlated with the duration of postIWI (i.e. subsequent sleep episodes) in the Lν_s_ phase only (Fig 6A). However the underlying correlations differed from those described above for preIWI. In the Lν_s_ phase, WAKE bout duration was positively correlated with postIWI cumulative time in NREM (Fig 6B) and mean NREM bout duration (Fig 7B), but not with postIWI bout count (Fig 7A). There were significant negative correlations between WAKE bout duration versus postIWI cumulative time in REM (Fig 6C) and postIWI bout count (Fig 7A) in the Lν_w_, Dν_w_ and Dν_s_ phases, along with a non-significant trend for negative correlation with mean postIWI REM bout duration (Fig 7C).

#### 3.5.2. NREM bout durations versus inter-NREM intervals (INI)

NREM bout duration was significantly positively correlated with the durations of preINI in the Lν_s_ phase only (Fig 6D). This was accompanied in Lν_s_ with positive correlations between NREM bout duration versus cumulative WAKE, mean WAKE duration (Fig 7E) and mean REM duration in preINI (Fig 7F). There were no correlations between NREM bout duration and preINI bout count in any circadian-ultradian phase (Fig 7D). There were significant positive correlations between NREM bout duration and preINI mean REM bout duration in all four circadian-ultradian phases (Fig 7F).

NREM bout duration was significantly positively correlated with the durations of postINI in the Lν_w_, Lν_s_ and Dν_s_ phases (Fig 6D). In all four circadian-ultradian phases there were significant positive correlations between NREM bout duration versus postINI cumulative REM (Fig 6F) and postINI mean REM bout duration (Fig 7F). There were weak, marginally significant negative correlations between NREM bout duration and postINI cumulative WAKE (Fig 6E) and postINI mean WAKE bout duration (Fig 7E) in the Lν_w_, Dν_w_ and Dν_s_ phases. Finally, there were significant positive correlations between NREM bout duration and postINI bout count in all four circadian-ultradian phases (Fig 7D).

#### 3.5.3. REM bout durations versus inter-REM intervals (IRI)

There was a significant positive correlation between REM bout duration and preIRI duration in the Lν_w_ and Lν_s_ phases (Fig 6G). This was accompanied by statistically significant positive correlations between REM bout duration versus preIRI cumulative NREM in the Lν_w_, Lν_s_ and Dν_s_ phases (Fig 6I) and preIRI mean NREM bout durations in all four circadian-ultradian phases (Fig 7I). These positive correlations were offset in the D phase by negative correlations between REM bout duration versus preIRI cumulative WAKE (Fig 6H) and preIRI bout count (Fig 7G).

REM bout duration was positively correlated with postIRI duration in the Lν_w_, Lν_s_ and Dν_s_ phases (Fig 6G), linked to corresponding positive correlations between REM bouts versus postIRI cumulative WAKE (Fig 6H) and cumulative NREM (Fig 6I). REM bout duration was positively correlated with postIRI bout count in Lν_w_ and Lν_s_, but not in the D phases (Fig 7G). REM bout duration was also positively correlated with postIRI mean NREM bout duration in all four circadian-ultradian phases (Fig 7I), and postIRI mean WAKE bout duration in the Lν_s_ and Dν_s_ phases (Fig 7H).

REM bouts are followed either by WAKE or NREM and this “choice” of post-REM trajectory is quantified by the relative probability of transition trajectory (P^*^_rn_ = 1-P^*^_rw_). During the Lν_s_ phase, short (i.e. sub-median) REM bouts were found to have significantly higher P^*^_rn_ than long (i.e. supra-median) REM bouts (P^*^_rn_ = 0.664 ± 0.066 versus 0.549 ± 0.119, paired t test P = 0.0206). A similar, but non-significant, trend was also present in the Lν_w_ phase, but was absent in the Dν_w_ and Dν_s_ phases. Thus, during the L phase of the circadian cycle, shorter REM bouts were followed by a greater proportion of postIRI’s beginning with NREM, whereas longer REM bouts were followed by a corresponding small majority of postIRI’s beginning with WAKE. Given the extremely low probability of WAKE to REM transitions (P_wr_, Fig 3A), REM onset very rarely began after WAKE and almost always occurred after NREM. As a consequence of this constraint, nearly all IRI’s beginning with NREM contained an odd number of bouts (R-N-[W-N]_*n*_-R; REM occured after 1, 3, 5… bouts) whereas IRI’s beginning with WAKE contained an even number of bouts (R-W-N-[W-N]_*n*_-R…; REM occured after 2, 4, 6… bouts) (Fig 8). IRI duration was strongly correlated with number of bouts (Spearman ρ; Lν_w_, 0.74 ± 0.09, Lν_s_, 0.65 ± 0.1, Dν_w_, 0.75 ± 0.08, Dν_s_, 0.73 ± 0.07; all P < 10^-7^). IRI’s starting with NREM consequently tended to be shorter than those beginning with WAKE.

**Fig. 8.**
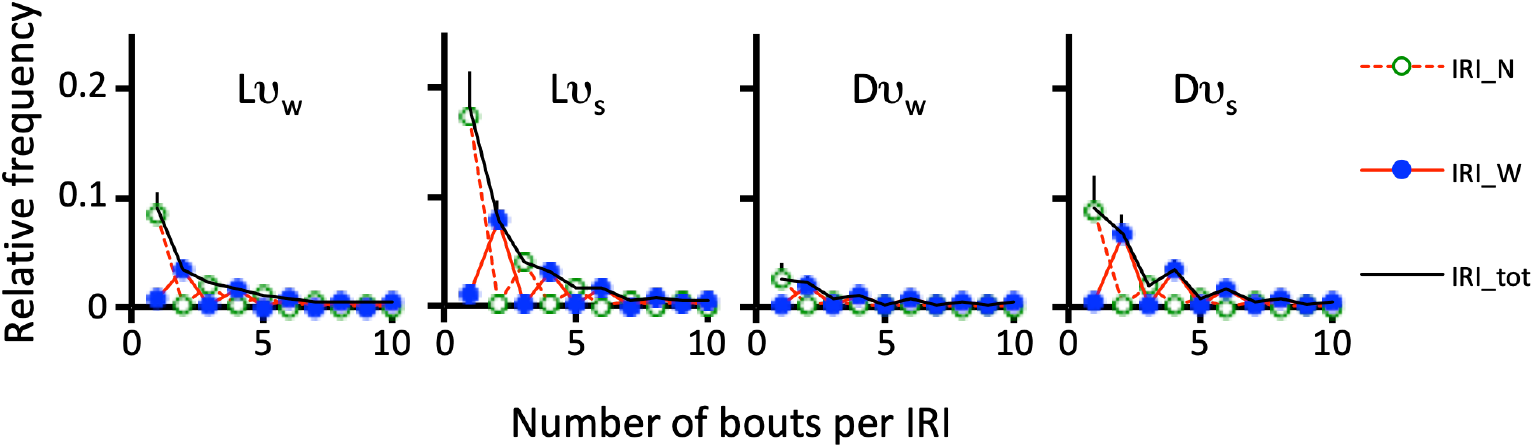
Relative frequency of inter-REM intervals (IRI) as a function of the number of bouts per IRI. To allow visualization of differences in overall frequencies across the four circadian-ultradian phases, relative frequency was calculated using the sum of all IRI’s over total recording time as the denominator. In each circadian-ultradian phase, shown in separate panels, the overall relative frequency curve (IRI_tot) is shown (mean + 95% CI, n = 10) in black. It does not describe a smooth geometric decline as would be predicted by a classical first-order Markov model. The IRI’s were subdivided according to whether they began with NREM (green circle, dotted red line; IRI_N) or with WAKE (blue symbol, solid red line; IRI_W). This shows that most IRI_N have an odd number of bouts and most IRI_W have an even number of bouts.

### 3.6. Sequences of sub-types of WAKE and sleep

Sleep episodes were categorized into two sub-types based on the number of bouts per episode - single-bout sleep episodes (S_s_) and multi-bout episodes (S_m_). Survival curves for each sleep sub-type are shown in Fig 1B and demonstrate that S_m_ episodes were distinctly longer than S_s_ episodes, as expected. This was the case in all four circadian-ultradian phases with the largest sub-type difference in Lν_s_ and the smallest difference in Dν_w_.

WAKE bouts were categorized into two sub-types on the basis of the bi-exponential survival curve (Fig 1A). A threshold duration (T_bl_) marked the transition from “brief” (W_b_) to “long” (W_l_) WAKE sub-types, and was defined on the basis of the point of intersection of the two linear segments of a semi-log plot (Fig 1A). Mean (± 95% CI) T_bl_ was Lν_w_, 2.25 ± 0.23 min; Lν_s_, 2 ± 0.5 min; Dν_w_, 2.5 ± 0.34 min, Dν_s_, 2.17 ± 0.49 min. Tukey’s multiple comparisons test determined that T_bl_ in Lν_s_ was significantly lower than that in Du_w_ (P_adj_ = 0.0246) but all other pairwise comparisons were non-significant.

χ^2^ tests were used to assess whether W_b_ and W_l_ sub-types were followed at random by S_s_ or S_m_ sleep sub-types, and vice-versa. Significant differences from random association were found for the WAKE to sleep sequences in the Lν_w_, Dν_w_, and Dν_s_ circadian-ultradian phases, but not in the Lν_s_ phase. In contrast, there were no significant deviations from random association in sleep to WAKE sequences in any circadian-ultradian phase.

The above result was examined in closer detail by calculating the percentage deviation of state sub-type sequences from the random order predicted by the null hypothesis. Fig 9A demonstrates that in the Lν_w_, Dν_w_, and Dν_s_ circadian-ultradian phases, the long W_l_ sub-type was 10 - 30 % more likely to be followed by S_s_, and 30 - 38 % less likely to be followed by S_m_ than would be expected on the basis of random chance. The same trend was also apparent in the Lν_s_ phase (Fig 9A), but this did not reach statistical significance.

**Fig. 9.**
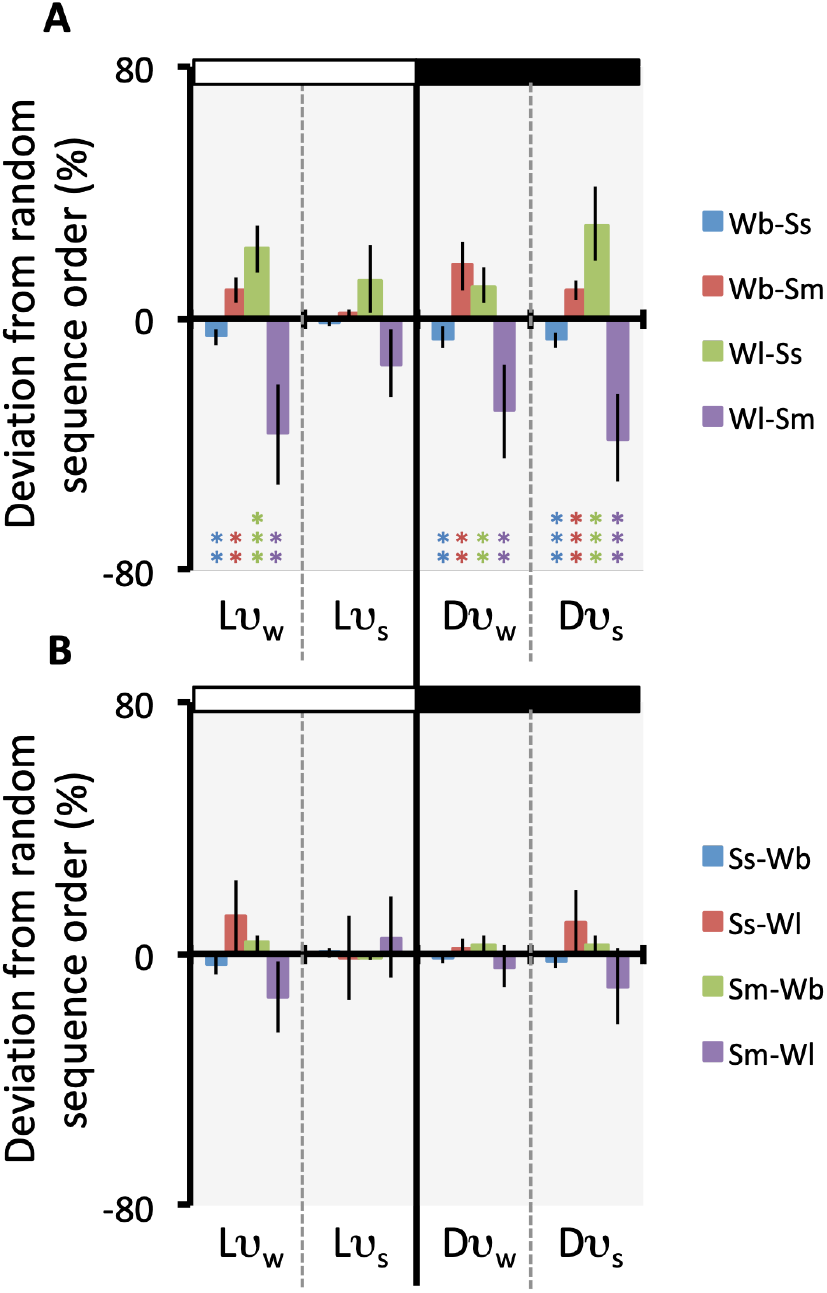
Sleep episodes were subdivided into those consisting of a single bout (S_s_) and those consisting of two or more bouts (“multi-bout sleep”, S_m_). WAKE bouts were also subdivided into “brief” (W_b_) and “long” (W_l_) sub-types (see Fig 1). It was of interest to know whether the sub-types of sleep were associated at random with the sub-types of WAKE. Using 2 x 2 contingency tables, the expected numbers of each sequence (null hypothesis of random association) were compared with observed numbers, and percent deviations (100 * observed / expected) were calculated. **A**. Percent deviation (mean ± 95% CI, n = 10) for WAKE to sleep sequences. Data show statistically significant non-random sequences; higher than expected occurrence of W_b_ to S_m_ and W_l_ to S_s_ sequences and correspondingly lower than expected occurrence of W_b_ to S_s_ and W_l_ to S_m_ sequences in all circadian-ultradian phases except Lν_s_. **B**. Percent deviation (mean ± 95% CI, n = 10) for sleep to WAKE sequences. Data show that both sleep sub-types were followed by the two WAKE sub-types in proportions that could not be distinguished from random association in all circadian-ultradian phases. One-sample t-test was used to determine statistical significance of group deviations: *, P < 0.05; **, P < 0.01; ***, P < 0.001; ****, P < 0.0001.

Furthermore, in the Lν_w_, Dν_w_, and Dν_s_ circadian-ultradian phases the short W_b_ sub-type was 9 - 16 % more likely than expected to be followed by S_m_ and approximately 6 % less likely than expected to be followed by S_s_. The W_b_ sub-type was followed in random proportions by S_s_ and S_m_ in the Lν_s_ phase (Fig 9A).

There were no statistically significant deviations from predicted random order in sequences of sleep to WAKE sub-types in any circadian-ultradian phase (Fig 9B).

## 4. Discussion

An unresolved question in sleep research concerns how longer-term rhythms interact with short-term sleep-wake patterns and, in turn, how all of these patterns arise within a single time-series of states, the durations and trajectories of which are determined by stochastic transitional events. This study was an attempt to shed light on this problem using statistical analysis of baseline recordings of rats. The focus was on the way state transition probabilities, both within and between sequences of states, vary as a function of joint circadian-ultradian phase.

Recent evidence suggests that longer-term patterns of sleep-wakefulness, such as diurnal rhythms and post sleep deprivation rebounds are mediated in part by adjustment of the relative durations of the ν_w_ and ν_s_ phases of the ultradian rhythm^16^. It was found that approximately 28% of the amplitude of the circadian rhythm in sleep and wakefulness was attributable to variation in the proportion of time spent in the ν_w_ and ν_s_ phases of the ultradian cycle (i.e. ultradian duty cycle, D_c_ = ν_w_ / [ν_w_ + ν_s_]); D_c_ was lower in L and higher in D. This implies that the remaining ∼72% of the circadian rhythm amplitude was attributable to L to D adjustment in state transition probabilities. The present study provides further insight into this issue by describing how state transition probabilities vary in relation to circadian and ultradian phase.

### 4.2. Long-term sleep-wake patterns: Variation in sleep-wake states and transition probabilities across circadian-ultradian phases

Fig 3A presents an overview of the mean values of state transition probabilities in relation to circadian and ultradian phases. Fig 3B takes the data in Fig 3A and isolates the ultradian phase differences in each of L and D, while Fig 3C isolates the circadian phase differences in each of ν_w_ and ν_s_.

During the L phase of the circadian cycle, there was a pronounced ultradian rhythm in P_wn_, and much smaller ultradian rhythms in P_nw_ and P_nr_. In the D phase, ultradian rhythms were observed in P_wn_, P_nw_ and P_rw_ (all three of comparable magnitude). Thus, the main factor determining the amplitude of the ultradian rhythm during the L phase was WAKE maintenance (P_ww_ ≈ 1 - P_wn_ when P_wn_ >> P_wr_; Fig 3A), whilst during the D phase, WAKE maintenance, and probabilities of arousal from NREM and REM were all prominently involved. This suggests that WAKE-related mechanisms predominate in the expression of ultradian sleep-WAKE rhythms in rats.

Since ultradian rhythms are superimposed upon, and therefore a component part of, circadian oscillations, any circadian modulation of transition probability must involve modulation of one or both phases of the ultradian oscillations, because both rhythms are composed of the same sequence of states. The results of the present analysis indicate that the amplitude of the circadian rhythm in sleep-wakefulness was influenced by mechanisms that modulate state transition probabilities in both the ν_w_ and ν_s_ phases of the ultradian cycles (Fig 3C). Furthermore, as was discussed above in the case of ultradian rhythms, WAKE-related transitions were also the main variables contributing to circadian amplitude.

In the ν_w_ phase of the ultradian cycles, P_wn_ was 53 % higher in L relative to D, and probabilities of arousal from NREM (P_nw_) and REM (P_rw_) were decreased by 54 % and 66 % respectively in Lν_w_ relative to Dν_w_ (Fig 3A,C). Thus, the extent to which WAKE was promoted in the WAKE-dominant phase of the ultradian cycle, ν_w_, was lower in L than in D and the net likelihood of arousal was lower in Lν_w_ versus Dν_w_. Similarly, in the ν_s_ phase of the ultradian cycles, P_wn_ was 45 % higher in L than in D (Fig 3A,C) and probabilities of arousal from NREM and REM were, respectively, 42 % and 52 % lower in Lν_s_ than in Dν_s_. These changes indicate an enhanced suppression of wakefulness during the Lν_s_ phase compared with the Dν_s_ phase.

The probabilities of NREM-REM cycling exhibited little or no circadian variation in both the ν_w_ and ν_s_ phases of the ultradian rhythm (Fig 3C). In the ν_w_ ultradian phase, P_nr_ and P_rn_ were non-significantly lower (by 19 % and 6 %, respectively) in L than in D. Likewise, in the ν_s_ ultradian phase, P_nr_ and P_rn_ were 21 % and 8 % lower in L than in D. This difference for P_nr_ (but not P_rn_) was marginally significant (Fig 3C).

In summary, the amplitude of the circadian rhythm in sleep and wakefulness was associated with primary modulation of WAKE-related bout-mean transition probabilities in both phases of the ongoing ultradian rhythm. Probability of WAKE maintenance (i.e. the additive inverse of P_wn_) was increased during both the ν_w_ and ν_s_ ultradian phases of the night (D), and suppressed during both ultradian phases in the daytime (L). These changes in WAKE maintenance probability, were accompanied by parallel changes in the probabilities of arousal from both NREM and REM.

### 4.2 Short term sleep-wake patterns: variation in transition probabilities within each state

The data presented in Fig 2 demonstrate that all transition probabilities varied with time in state. The bold coloured lines indicate the between-animal means and the light lines indicate the curves for each individual animal. It is clear that there was substantial quantitative variability between animals, but the overall timecourse was nevertheless fairly consistent. The increasing variability with increasing time in state is a reflection of the progressively decreasing precision of this approach to probability estimation; fewer bouts remain in the sample as duration of bout increases. Furthermore, whilst there were clear quantitative differences between probability profiles across circadian-ultradian phases, both in terms of average probability levels as well as rates of change with time in state, it is noteworthy that the overall shape of the probability profiles were reasonably consistent across phases. This result confirms a prior study of REM sleep over circadian time^10^.

### 4.3. Short term sleep-wake patterns: serial correlations between contiguous states

This study confirms the presence of statistical dependence between adjacent states. With a few minor exceptions, short-term dependence was qualitatively and quantitatively similar in the four circadian-ultradian phases (Fig 4). These data confirm that durations of state bouts are influenced by prior state^12^. McShane and colleagues studied this issue in several strains of mice using a “spike and slab” analysis of bout durations and noted differences between strains, indicating variability in underlying mechanisms. They did not take ultradian phase into account and a direct comparison with rat data in the present study is not possible. Nevertheless, in a broad sense the data are consistent in that categorical associations in both studies point to 2nd-order transition dynamics in both species. In the present study, the most prominent correlations were observed in state pair sequences involving REM. Specifically, there were statistically significant positive correlations in REM to WAKE pairs, in REM to NREM pairs and in NREM to REM pairs. This suggests that neural mechanisms involved in the control of REM play a major role in the short-term statistical patterning of overall sleep-wake sequences^10^.

Positive serial correlations imply a non-random association of bout durations in contiguous bouts of different states. It was also found that this association extended further, as indicated by significant serial correlations between sequential bouts within a given state (Fig 5). This was the case for WAKE bouts, NREM bouts and sleep episodes (which consist of a variable number of bouts), but not for REM bouts. The latter result may be explained in part by greater separation between REM bouts in terms of both time elapsed and number of bouts per IBI, compared with the other states.

### 4.4. Short term sleep-wake patterns: serial correlations between bouts and contiguous inter-bout intervals

In Fig 6A,D,G the Spearman rank correlation coefficient (ρ) is shown for bout duration versus prior (“pre”) and subsequent (“post”) inter-bout intervals for all three states in each of the four circadian-ultradian phases. Attention is drawn to the following prominent features of the results: (i) correlations were weak for WAKE versus IWI (sleep), intermediate for NREM versus INI and strongest for REM versus IRI, (ii) within each state, ρ values varied across circadian-ultradian phases, with generally stronger correlations in the sleep-dominant L phase of the circadian cycle and ν_s_ phase of the ultradian cycle, and (iii) the bout versus postIBI correlations tended to be stronger than the bout versus preIBI correlations.

Further analysis found that the bout duration versus IBI correlations were the net result of correlations between bouts and different components of the IBI intervals. Each IBI is composed of two states, and this study has confirmed previous reports that the bout - IBI correlations were related to IBI cumulative state content (Fig 6)^8, 13, 15^. As mentioned, there were instances of significant correlations of opposing sign that negated each other, sometimes resulting in non-significant net bout - IBI duration relations and these interactions are discussed separately for each state below.

#### 4.4.1. WAKE and inter-WAKE intervals (IWI, sleep episodes)

Overall, WAKE bout durations were only weakly correlated with durations and composition of prior and subsequent sleep episodes (Figs 6A-C, 7A-C). WAKE bout durations tended to be positively correlated with preIWI REM bout duration, preIWI cumulative REM content and, to a small extent, preIWI bout count. Conversely, WAKE bout durations tended to be negatively correlated with postIWI cumulative REM, and postIWI bout count, with a non-significant trend for negative correlation with mean postIWI REM bout duration. The net positive correlation between WAKE bout duration and postIWI duration in Lν_s_ was associated with positive correlations of WAKE bout duration with mean postIWI NREM bout duration and postIWI cumulative NREM in that phase only. The WAKE – postIWI correlations arose primarily from a reorganization of state transition trajectory (thereby affecting bout count) in the Lν_w_, Dν_w_, and Dν_s_ circadian-ulradian phases and to mean NREM bout duration in the Lν_s_ phase only. The weak WAKE - preIWI correlation was mainly linked to preIWI mean REM bout duration, which corresponds to the REM-WAKE correlation shown in Fig 4.

Collectively, these observations have relevance to some basic tenets of contemporary sleep theory. It is widely assumed that a putative “sleep homeostat” is active under baseline conditions to a degree that varies with circadian phase^26^. This hypothesis posits that sleep duration is proportional to the duration of prior wakefulness^27^; that is, a positive serial correlation should be observable between WAKE bout durations and contiguous sleep episode durations. Mistlberger and colleagues^28^ examined this prediction in circadian arrhythmic rats (after lesions of the suprachiasmatic nuclei) and reported weak positive correlations for both WAKE to sleep and sleep to WAKE sequences, although the data selection protocols used in the analysis may have been compromised by failure to account for ultradian rhythms, and very few animals were considered to be statistically significant. A similar result was obtained by Webb and Friedmann^29^, who used a χ^2^ test to assess defined classes (short, medium and long) of bout duration in rats – any correlations, if present, were weak and not statistically significant by this procedure. The latter study also did not account for ultradian phase in the analysis.

The present study has confirmed the absence of significant positive serial correlations between WAKE and subsequent sleep durations in the Lν_w_, Dν_w_, and Dν_s_ circadian-ultradian phases, and found a very weak and marginally significant correlation in the Lν_s_ phase only (Fig 6A). These results, together with those published recently^16^, are not supportive of a sleep homeostat hypothesis.

This issue was examined further in the present study by modifying the approach used by Webb and Friedmann^29^. Here, both states were partitioned into two sub-types; WAKE was divided into “brief” (W_b_) and “long” (W_l_) sub-types using a threshold duration based on the bi-exponential survival curve of WAKE bouts^6^. Sleep episodes, which could not be partitioned by the same threshold-based method, were divided into single-bout (S_s_) and multi-bout (S_m_) sub-types, which differ substantially in their episode duration distributions (Fig 1). It was found (Fig 9A) that in the Lν_w_, Dν_w_, and Dν_s_ circadian-ultradian phases, the frequencies with which the two WAKE sub-types were followed by each of the two sleep sub-types was significantly non-random. That is, in comparison with the numbers expected by random chance, long W_l_ bouts were followed by more of the shorter S_s_ and less of the longer S_m_, and the converse was true for W_b_ bouts. This is the opposite of the sleep homeostat prediction. The alternate sequence - sleep sub-type to WAKE sub-type - was found to be randomly organized (Fig 9B). It is therefore concluded on the basis of these results that sleep episode duration is not regulated in proportion to prior WAKE bout duration by a short-term sleep homeostat mechanism in the rat.

#### 4.4.2. NREM and inter-NREM intervals (INI)

NREM bout duration was significantly positively correlated with preINI duration in the Lν_s_ phase only, linked to a positive correlation with mean preINI WAKE bout duration (Fig 7E). This corroborates the positive WAKE - NREM correlation observed only in Lν_s_ shown in Fig 4. There were pronounced positive correlations between NREM bout duration and preINI mean REM bout duration in all circadian-ultradian phases (Fig 7F), again corroborating the result from paired states (Fig 4).

NREM bout durations were significantly positively correlated with postINI duration in the Lν_w_, Lν_s_ and Dν_s_ phases (Fig 6D), linked to positive correlations (in all four circadian-ultradian phases) with postINI bout count (Fig 7D) and postINI mean REM bout duration (Fig 7F). These positive associations were opposed in the Dν_w_ phase by a negative correlation between NREM bout duration and postINI mean WAKE bout duration (Fig 7E).

The above results indicate that the durations of NREM and both pre- and post-INI are linked via the serial correlations between NREM and REM (i.e. rN and nR in Fig 4). The correlation between NREM bout duration and postINI bout count (Fig 7D) was also indirectly dependent on REM because NREM - INI dynamics were strongly constrained by the very low probability of sleep-onset REM (P_wr_). Specifically, the very low P_wr_ restricted the number of bouts per INI so that, except on very rare occasions, only three sequences were seen: NREM-REM-NREM (single-bout INI), NREM-WAKE-NREM (single-bout INI), and NREM-REM-WAKE-NREM (dual-bout INI). It was found that in all four circadian-ultradian phases, long (supra-median) NREM bouts were significantly more likely than short (sub-median) NREM bouts to transition to REM (i.e. P^*^_nr [long]_ > P^*^_nr [short]_; paired t test, all circadian-ultradian phases P ≤ 0.0015). Furthermore, Fig 4 demonstrates that in all four circadian-ultradian phases there was a significant positive correlation between NREM bout duration and subsequent REM bout duration. Thus, there was a bias for relatively long NREM bouts to transition to REM and for these REM bouts to also be relatively long. Taking this further, it was also found that in the L phase, long (supra-median) REM bouts were more likely than short (sub-median) REM bouts to transition to WAKE (i.e. P^*^_rw [long]_ > P^*^_rw [short]_) but this was statistically significant only in the Lν_s_ circadian-ultradian phase (paired t test, P = 0.02). Finally, there was a positive correlation between REM bout duration and subsequent WAKE duration (Fig 4) so that longer REM bouts were more likely to be followed by relatively long WAKE bouts.

Collectively, these serial correlations indicate that there was a tendency (albeit weak because the correlation coefficients were weak to moderate) for longer NREM bouts to be followed by longer dual-bout INI’s with relatively long REM and WAKE bouts, and shorter NREM bouts to be followed by shorter single-bout INI’s consisting of a relatively short REM or WAKE bout. This underscores the inter-dependence of the different state transition mechanisms and provides a good example of how the dynamics of one state (in this case NREM) can be strongly influenced by mechanisms that control transitions between the other two states (in this case P_wr_)^19^.

#### 4.4.3. REM and inter-REM intervals (IRI)

Kripke and colleagues were the first to report a weak positive correlation between REM bout duration and subsequent inter-REM interval (postIRI) in rhesus monkeys (*Macaca mulatta*)^30^. Ursin^15^ subsequently observed the same in cats and proposed that the REM – postIRI correlation could be due to a preparatory mechanism in which REM sleep is permissive of NREM expression. This hypothesis has since been widely accepted and is often referred to as a short-term REM homeostat^8, 31, 32^.

Vivaldi and colleagues, working on rats, have contributed the most detailed studies relating to REM – IRI correlations^8, 10, 13, 14^. The present results have corroborated many of their findings, and extended them through an examination of ultradian phase. They reported a correlation between REM bout duration and postIRI duration in both light and dark phases of the diurnal cycle^13^ and the present study extends that conclusion to both of the ultradian phases in L. However, it was found here that the correlation was absent in the Dν_w_ circadian-ultradian phase, likely due to the low number of REM - IRI cycles that occur in that phase (Fig 8), coupled to the high variability of IRI durations related to very variable WAKE bout durations. Despite this, positive correlations were observed between REM bout duration and postIRI mean NREM bout duration in all four circadian-ultradian phases (Fig 7I). Positive correlation with postIRI bout count (Fig 7G) were observed in the L phase only and this helps explain the stronger overall REM bout - postIRI duration correlation in the L phase ^13^.

One interesting result in the present study, only rarely reported in previous studies^33^, was a significant REM – preIRI correlation in the L circadian phase only (Fig 6G). This result is consistent with a short-term REM homeostat but it is not clear why such a mechanism would be absent at night. The present data suggest a possible explanation for this L versus D discrepancy. It was found that there was a moderately strong correlation between REM bout duration and preIRI mean NREM bout duration (Fig 7I), which concurs with the positive serial correlation between the durations of contiguous NREM and REM bouts (nR sequence pairs; Fig 4). However, although these positive correlations were observed in all four circadian-ultradian phases, in the D phase they were offset by negative correlations between REM bout duration and preIRI bout count (Fig 7G), preIRI cumulative WAKE (Fig 6H) and (non-significantly) preIRI mean WAKE bout durations (Fig 7H). These results again highlight the complex co-dependence of states in overall pattern generation^19^.

The implications of the near absence of WAKE to REM transitions (P_wr_; SOREM onset) for the inter-NREM interval has already been discussed and here it is argued that it has an equally important impact on IRI dynamics. Negligible P_wr_ constrains the patterns of state cycling that are possible - in addition to reducing the frequency of REM bouts, very low P_wr_ also has implications for sequences of all three states. The inter-REM intervals (IRI) consist of alternating NREM and WAKE bouts. Each IRI can *begin* with transition from REM to either NREM or WAKE, but because of the negligible P_wr_, each IRI nearly always *ends* with a transition from NREM to REM. As a consequence of this constraint, IRI’s that begin with NREM nearly always have an odd number of bouts (REM-NREM-[WAKE-NREM]_n_-REM) and IRI’s that begin with WAKE nearly always have an even number of bouts (REM-[WAKE-NREM]_n_-REM). IRI duration is strongly correlated with number of bouts, and the percentage of IRI’s declines approximately geometrically with increasing bout number (Fig 8). Consequently, IRI’s beginning with NREM tend to be shorter than those beginning with WAKE, owing to the relatively large number of single-bout IRI’s (i.e. REM-NREM-REM sequences). Thus, the relative frequencies of REM arousals versus REM to NREM transitions has an influence on IRI duration as a direct consequence of the strongly attenuated P_wr_.

It was found in the present study that state transition probabilities change over time in state, as shown previously for REM^10^. In the case of REM, P_rw_ (probability of arousal) and P_rn_ (probability of REM to NREM transition) differed in their timecourse across a REM bout. Specifically, during the L circadian phase, at the start of a REM bout P_rn_ was approximately double that of P_rw_, yielding a relatively high probability of NREM onset (P^*^_rn_ approximately 0.67). This difference between transition probabilities then progressively decreased until, after about 2 minutes of REM, P_rw_ and P_rn_ were approximately equal (P^*^_rn_ ≈ 0.5). Hence, shorter REM bouts tended to be followed by shorter IRI’s in part because they have a higher tendency to begin the IRI with NREM, and therefore tended to have fewer bouts in postIRI.

### 4.5. Implications for models of sleep-wake patterns

Attempts to model sleep-wake patterns have used a variety of quantitative approaches, the overall objective being to develop a conceptual framework with which to interpret and predict normal and abnormal sleep-wake patterns under different circumstances, and to help understand how they are controlled^34, 35^.

Rodent and human sleep-wake states have been modeled as probabilistic systems exhibiting the Markov property of first order dependence, where state transitions are “memoryless” in the sense that that they occur with a probability that is conditional on current state and independent of prior states^1-4, 6, 23, 36-42^. Hence, in a classical Markov model, conditional probability of state maintenance and relative probability of transition trajectory, are assumed to remain constant with time in state, giving rise to exponential state bout survival curves. However, the present study concurs with prior studies^9-11^ by showing that transition probabilities did vary over time in state, with a pattern that was consistent across animals.

As discussed above, the present analysis also identified serial correlations in bout durations between consecutive states (Fig 4) as well as between successive bouts within the same state (WAKE and NREM, but not REM; Fig 5). This implies a bout-to-bout statistical dependence that extends over sequences of at least 2 or 3 bouts and implies a higher-order system^6^ where prior state-time history can influence current transition probabilities. Overall, these data indicate that state transition probabilities vary both during and across states.

Whilst debate has revolved around how a presumptive short-term REM homeostat can produce the positive REM – postIRI correlation, the possibility of an alternative (non-homeostatic) explanation for the correlation has received much less attention^43, 44^.

A new hypothesis for the statistical origin of higher-order associations within the rat hypnogram, including the REM-postIRI correlation, can be proposed based on the observations of within-bout variation in transition probability together with significant serial correlations spanning 2 – 3 bouts. The serial Spearman rank correlation coefficient is used in this context as an autocorrelation function describing the normalized covariance of ranked durations in defined pairs of state bouts. As such, it quantifies the conditional bout-to-bout persistence or “momentum” in state stability. Hence, the positive correlation coefficients identified in the present study are consistent with the hypothesis that the state transition probability at the end of a bout may be carried over to some extent to the beginning of the next bout.

If total transition probability (P_im_) can be taken as a measure of state instability, then the profiles shown in Fig 2 indicate that in all three states, and in all four circadian-ultradian phases, each state bout begins with a brief interval (approximately 1 - 5 min) of steady or decreasing instability, gradually turning over to a progressive increase. This indicates that the state transition system does not conform to a classical Markov process and that a semi-Markov model is more appropriate, as was suggested by Yang and Hursch half a century ago^45^. Furthermore, serial correlations between contiguous states imply that the system is not 1st-order, and higher order dependence was incorporated into a computer model as a “transition momentum” function. This model (termed “SM3m”) is a 3-state 2nd-order semi-Markov model and is shown schematically in Fig 10.

**Fig. 10.**
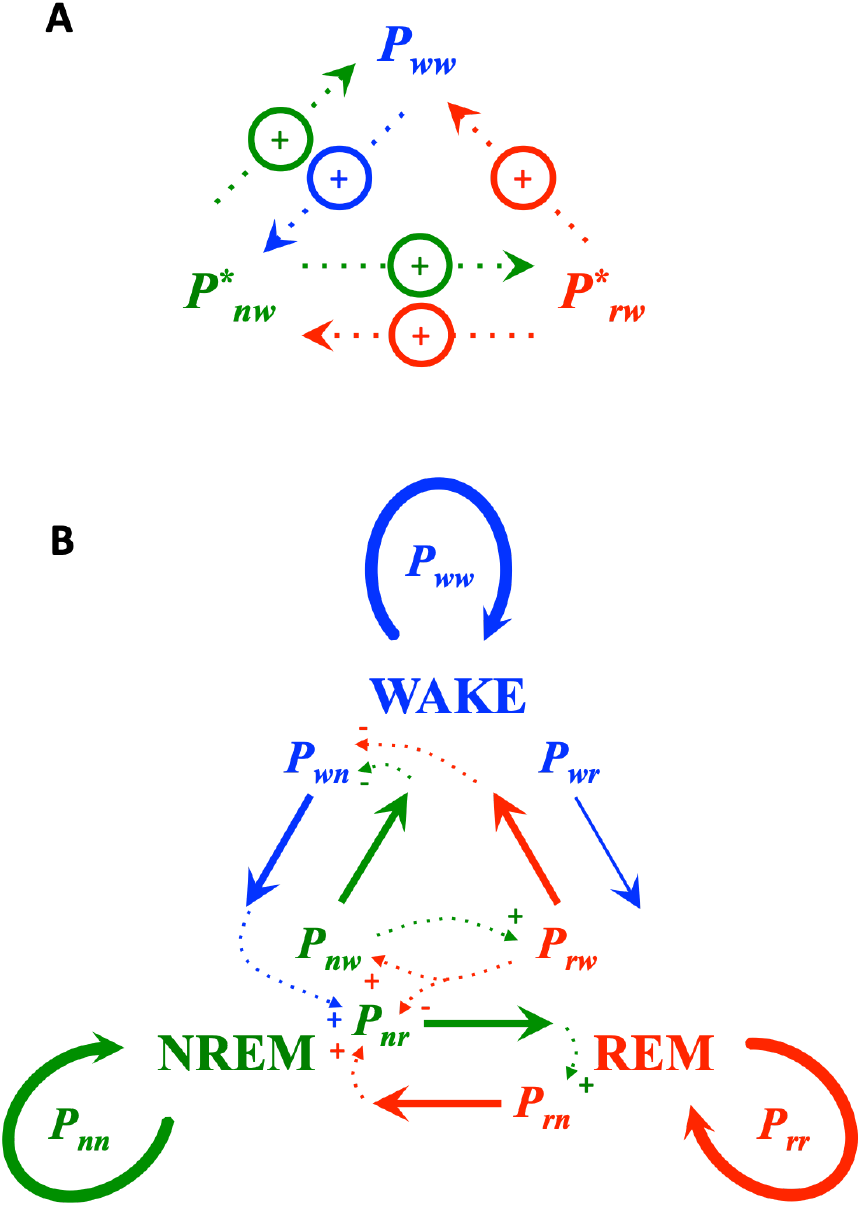
Schematic of the 2nd-order three-state semi-Markov model (SM3m) used to simulate short-term sleep-wake patterns. Each solid arrow indicates conditional probability of state transition, quantified in this paper on a per epoch (5s) basis. Inter-state momentum of transition probability is indicated by coloured dotted arrows. Here, probability of state transition at the end of a bout influences the indicated state transition probability at the start of the next bout. Signs indicate augmentation (+) and attenuation (-). A. Schematic explanation of the basic concept behind the SM3m model, which was designed as a Markov chain with 2nd-order “WAKE momentum”. Here, high wakefulness “drive” during WAKE is retained to some degree in the next sleep episode and manifests as an elevated relative probability of arousal. Likewise, a high probability of arousal during sleep is retained to some extent as elevated WAKE maintenance after awakening. High relative probability of arousal during NREM is retained to some degree during REM and vice-versa. The gain of momentum was adjustable in simulations. B. Each state is subject to three transition probabilities shown as solid coloured lines (WAKE, blue, NREM, green, REM, red). It was convenient to compute momentum via absolute probabilities of state maintenance (P_ww_, P_nn_, P_rr_) and relative probabilities of arousal (P^*^_nw_, P^*^_rw_), and to then use these primary variables to calculate absolute post-bout transition probabilities (P_wn_, P_nw_, P_nr_, P_rw_, P_rn_). SOREM’s were disallowed (P_wr_ = 0), so all sleep onsets occurred via NREM (and there is therefore no relative transition probability for post-WAKE trajectory). P_rr_ was assumed to remain unaffected by “momentum” so that variations in P^*^ reflect symmetrical variation of absolute transition probabilites (ΔP_rw_ = ΔP_rn_). P_nn_ was assumed to vary in proportion to P^*^_nw_ so that variations in P^*^_nw_ reflect asymmetric variation in absolute transition probabilities (ΔP_nw_ < ΔP_nr_). Dotted lines indicate momentum pathways of absolute transition probabilities.

Full description, validation and sensitivity analysis of the SM3m model is beyond the scope of the present paper and only a few important results of quantitative model simulations are summarized briefly here. SM3m was able to reproduce the key features described in rat data, including REM – postIRI correlations, non-random sequential pairings of WAKE-sleep sub-type pairs and serial correlation between contiguous states.

The SM3m model shown in Fig 10 falls into the general class of “autonomous toy models”^46^. The two-process model^26^ and the Lotka-Volterra model^47^ are two other prominent models of this type. Such models are intentionally over-simplified and idealized, and for that reason they cannot be used to show that a hypothesis is a valid description of reality - the best that they can do is to establish plausibility^46^. Computer simulations using the SM3m model have confirmed the plausibility of the notion that state instability, quantified by transition probability, might be a state-spanning phenomenon, but a model cannot be accepted on the basis of plausibility alone.

In light of the above, the observation of a positive correlation between REM bout duration and subsequent IRI duration can now potentially be “explained” by two very different putative mechanisms: the REM homeostat^8^ and the inter-state transition momentum hypotheses. Both are plausible models but unfortunately both are also weak inferences and cannot be distinguished by using the approach used here for the simple reason that correlational data cannot infer causal mechanisms. The REM – postIRI correlation is consistent with both proposed mechanisms and therefore cannot be considered to represent “support” for either; both hypotheses are underdetermined. This study presents only indirect evidence pertaining to the new “inter-state transition momentum” hypothesis and direct experimental interventions are required to test the idea. The same criticism applies to the putative REM homeostat. The inter-state momentum hypothesis has the advantage that it specifies an identifiable mechanism that explains the full complement of statistical patterns, whereas the state transitional events underlying a REM homeostat remain opaque and it has not yet been shown to explain many of the observed statistical features of sleep-wake patterning.

The analysis used in this study rests on the assumption that the mechanisms controlling sleep-wake state expression act primarily by modulating the probability of state transition events. Variability of state bout durations and the unpredictability of the precise timing of transitions are prominent and ubiquitous features of the mammalian sleep-wake system, and full quantification of probabilities of state transition can have advantages when combined with conventional measures such as bout duration, bout frequency and cumulative time in state^48^. For example, quantitative deviations in statistical characteristics of sleep-wake state can be used as a sensitive approach to distinguish normal (e.g. “consolidated”) and disordered (e.g. “fragmented”) conditions^42, 49^.

Probabilistic models emphasize the significance of stochastic dynamics, and this begs the question as to why the state control system works this way, and whether stochastic state fluctuations reflect an advantageous property or unavoidable by-product of system design. Stochastic “noise” can be surprisingly beneficial for system performance in certain circumstances^50^ and is a potentially interesting area of theoretical and experimental investigation for sleep-wake regulation. Whilst deterministic models place emphasis on average patterns, stochastic models focus attention on the central role of dynamic variability, and encourage research in understanding its origin, regulation and functional significance. The unpredictability of transition events, together with the high proportion of very brief bouts, also raises interesting questions about the types of functional processes that might be dependent on sleep. Putative sleep functions must be compatible with frequent and unpredictable spontaneous interruption of sleep in polyphasic rodents.

### 4.6. Critique

This study has attempted to shed some light on sleep-wake patterns in rats. However the approach used here can be questioned regarding the relevance and generalizability of the data. The rats were held in “standard” laboratory conditions and recorded via implantable biotelemetry devices in the hope that minimal disturbance would yield reduced variability and thereby facilitate identification of “baseline” patterns of state expression. These recording conditions might be standard insofar as sleep laboratories are concerned, but they are far from typical for this species living under more natural conditions. The rats used here were all male (to avoid having to account for oestrus cycles in females), confined to a small cage and living in partial isolation (no visual or tactile contact with other rats, but auditory and olfactory stimuli were present), with minimal environmental stressors. Light cycles were constant (no variation due to cloud cover or lunar cycles for example), thermal fluctuations were minimized, wind and rain were absent, the room was soundproofed, humidity was held constant, food and water were readily available at all times, the rats were not able to exercise significantly (e.g., no exploratory or foraging acitivities), no conspecific competition or other interactions, no fighting, courtship, play, or threat of predation, no disease or parasites. All of these variables (and others not mentioned) that might be expected to be a normal part of the daily life of a rat were absent during recordings. These factors would be expected to influence wakefulness and sleep drives and therefore substantially influence the probabilities of state transition. It is therefore reasonable to suspect that the results reported herein might be atypical and that different conclusions might be anticipated in animals living under natural conditions. Advances have been made in this area in this area in recent years^51-57^ and it is clear that such studies would benefit greatly from the application of detailed studies of state transitions to better understand sleep-wake dynamics of free-living animals.

The present study uses data that were not collected specifically for this analysis, and in some respects were suboptimal. A major objective was to determine the reciprocal relations between state transition probabilities and circadian-ultradian phase. Hence the data in each animal were subdivided into four circadian-ultradian phase categories, which were pooled across the week of recordings. This analytical approach assumes that long-term variation was absent or at least randomized (i.e. no systematic trends) but this assumption was not tested and may not necessarily be valid. Another assumption built into the analysis, owing to the adoption of a binary scoring system for circadian and ultradian phases, was that state transition probabilities are unchanging as a function of time within each phase. This is unlikely to be true^6, 10^. Further subdivision of the circadian and ultradian cycles would be preferable, but this would necessitate a commensurate increase in sample sizes. For example, a simple subdivision of both circadian and ultradian rhythms into two parts (early-late L, early-late D, early-late ν_w_ and early-late ν_s_) would require a month of data just to retain the same statistical power. Thus, future studies of this kind will need to plan for very long-term recordings.

Another problem related to sample sizes relates to the use of a frequentist method for the estimation of transition probability. As mentioned earlier, sample sizes must be as large as possible if the estimate of probability obtained using ratios of bouts is to be a reliable indicator of the true probability of an underlying stochastic process. When calculating intra-bout probability profiles, the sample size progressively decreases with increasing bout duration and so the precision (and reliability) of the estimate of probability decreases proportionally. This means that only the early part of the state transition profiles can be used with confidence in model parameter assignment.

A related issue concerns the assumption of homogeneity of the sample. This assumption is implicit in this approach. As mentioned, probability is calculated as relative frequency in a sample of bouts that were pooled according to circadian-ultradian phase. This implicitly assumes that all bouts in a sample have the same time-in-state probability profile, but this is unlikely to be true. As already discussed, it is unlikely that circadian and ultradian phases are homogeneous, but the data also indicate that on a shorter time scale there are bout-to-bout autocorrelations that further imply a lack of sample homogeniety. The extent to which the data and conclusions of the present study may be affected by this issue is unknown, but the individual animal bout probability timecourses shown in Fig 2 most likely represent averages across a range of individual bout probability profiles. The SM3m model explicitly proposes that the profiles are indeed quite variable from bout to bout. Further research and refinement of the model, using analytical methods that do not involve averaging across bouts, will be required to shed light on these issues.

### 4.7. Concluding comments

Circadian and ultradian rhythms are primarily controlled by mechanisms that modulate WAKE-related transition probabilities. The data indicate that transition probabilities change with time in state, suggesting a semi-Markov process, and significant serial correlations between durations of contiguous states suggest state-spanning processes in a 2nd or higher order system. The data cast further doubt on the validity of the hypothesis that sleep is regulated in proportion to prior wakefulness, and present a plausible alternative to the hypothesis that REM sleep is regulated by a short-term homeostatic mechanism.

## Acknowledgements

Data used in this study was collected with the support of a Discovery Grant from the Natural Sciences and Engineering Research Council of Canada. The re-analysis of data and preparation of this paper were unfunded.

